# Destabilization of neuromuscular junctions and deregulation of activity-dependent signalling pathways in Myotonic Dystrophy type I

**DOI:** 10.1101/2023.03.08.531688

**Authors:** Denis Falcetta, Sandrine Quirim, Ilaria Cocchiararo, Mélanie Cornut, Marine Théodore, Adeline Stiefvater, Shuo Lin, Lionel Tintignac, Robert Ivanek, Jochen Kinter, Markus A. Rüegg, Michael Sinnreich, Perrine Castets

## Abstract

Myotonic Dystrophy type I (DM1) is the most common muscular dystrophy in adults. Previous reports have highlighted that neuromuscular junctions (NMJs) deteriorate in skeletal muscle from DM1 patients and mouse models thereof. However, the underlying pathomechanisms and their contribution to muscle dysfunction remain unknown. We compared changes in NMJs and activity-dependent signalling pathways in *HSA^LR^* and *Mbnl1^ΔE3/ΔE3^* mice, two established mouse models for DM1. DM1 muscle showed major deregulation of calcium/calmodulin-dependent protein kinases II (CaMKIIs), which are key activity sensors regulating synaptic gene expression and acetylcholine receptor (AChR) recycling at the NMJ. Both mouse models displayed increased fragmentation of the endplate, which preceded muscle degeneration. Endplate fragmentation was not accompanied by changes in AChR turnover at the NMJ. However, expression of synaptic genes was up-regulated in DM1 muscle, which may be linked to the abnormally high activity of histone deacetylase 4 (HDAC4), a known target of CaMKII. Consistently, expression of myosin heavy chains was deregulated as well, leading to a major switch to type IIA fibres in *Mbnl1^ΔE3/ΔE3^* muscle, and to a lesser extent in *HSA^LR^* muscle. Interestingly, although HDAC4 was efficiently induced upon nerve injury, synaptic gene up-regulation was abrogated in DM1 muscle, together with a reduced increase in AChR turnover. This suggested that HDAC4-independent mechanisms lead to the defective response to denervation in DM1 muscle. Our study shows that activity-dependent signalling pathways are disturbed in DM1 muscle, which may contribute to NMJ destabilization and muscle dysfunction in DM1 patients.

## INTRODUCTION

Myotonic Dystrophy type I (DM1) is a multisystemic disorder caused by expanded CTG triplet repeats in the 3’UTR of the *DMPK* (*Dystrophia Myotonica Protein Kinase*) gene, leading to muscle wasting, weakness and inability to relax (myotonia). Accumulation of toxic transcripts containing the expanded CUG repeats lead to the nuclear sequestration of splicing factors. The consecutive mis-splicing of specific genes is determinant in the pathogenesis of DM1-associated muscle alterations.^1, 2^ For example, mis-splicing of the *CLCN1* gene encoding the chloride channel ClC-1 is implicated in myotonia development. Mis-splicing of several genes encoding proteins of Ca^2+^-associated signalling pathways has also been shown to contribute to muscle dysfunction. Especially, previous reports unveiled that mis-splicing of *CAMK2* genes, encoding Ca^2+^/calmodulin-dependent protein kinases II (CaMKIIs), is a hallmark of DM1.^3–5^ However, the functional consequences of CAMKII deregulation in DM1 muscle and its contribution to DM1 pathogenesis were not explored.

CaMKIIs are important for the maintenance of neuromuscular junctions (NMJs), which are the synapses connecting motor neurons to muscle fibres. Especially, CaMKIIs enhance the recycling of acetylcholine receptors (AChRs) upon their internalization in the sub-synaptic region of muscle fibres (*i.e.* the endplate).^6^ Moreover, CaMKIIs indirectly repress the expression of synaptic genes in non-synaptic regions of innervated muscle, by inhibiting the myogenic transcription factor myogenin and the histone deacetylase 4 (HDAC4).^7–9^ Hence, deregulation of CaMKII signalling pathway may affect NMJ maintenance, by perturbing the expression and the dynamics of synaptic proteins.

Early studies on DM1 muscle biopsies pointed to NMJ-associated abnormalities, such as a reduced number of presynaptic vesicles, enlarged endplates or angular muscle fibres.^10–13^ Lack of denervation markers (*e.g.,* non-junctional AChR clusters) and the absence of massive motor neuron loss rejected the hypothesis that spontaneous denervation is part of DM1 pathogenesis.^14, 15^ NMJ alterations have also been reported in *DMSXL* mice and in mice deficient for Muscleblind-like 1/2 (Mbnl1/2), two mouse models for DM1, as well as in C. *elegans* DM1 mutants.^16, 17^ Moreover, nuclear foci, characteristic of DM1-associated accumulation of toxic RNA, were detected at the endplate, which may alter the expression of synaptic genes.^18^ Mis-splicing of some synaptic genes has been reported in muscle cells from DM1 patients.^19^ Hence, NMJ deterioration in DM1 is likely to contribute to muscle dysfunction, but the underlying pathomechanisms remain unknown.

Here, we analysed changes in activity-dependent signalling pathways in *HSA^LR^* and *Mbnl1^ΔE3/ΔE3^* mice, two established DM1 mouse models. Both mouse models displayed increased fragmentation of the endplate, which preceded muscle alterations. *HSA^LR^* and *Mbnl1^ΔE3/ΔE3^* mice also showed deregulation of CaMKII / HDAC4 signalling in muscle, which was accompanied by an increased expression of synaptic genes and a switch to slower fibre type. Interestingly, endplate remodelling upon nerve injury was hampered in DM1 mice, despite efficient induction of HDAC4. Hence, perturbations in activity-dependent signalling pathways may contribute to NMJ destabilization and muscle dysfunction in DM1 patients.

## RESULTS

### DM1-associated muscle alterations are similar in *Mbnl1^ΔE3/ΔE3^* and *HSA^LR^* mice

To evaluate changes in NMJs and activity-dependent pathways in DM1, we selected the *Mbnl1^ΔE3/ΔE3^* and *HSA^LR^* mouse lines, which are two well-established DM1 mouse models. In *Mbnl1^ΔE3/ΔE3^* mice, deletion of *Mbnl1* exon 3 leads to ubiquitous depletion of the splicing factor MBNL1 (*i.e.,* including in skeletal muscle and motor neurons^).21^ In contrast, *HSA^LR^* mice express the *HSA* transcript (*Human Skeletal Actin*) with long (CTG) repeats only in skeletal muscle, which allowed to unveil cell-autonomous defects.^20^ First, we compared the muscle phenotype between the two mouse models. There was no dystrophic sign in muscles from 3-months-old mutant mice (*Figure* 1A and 1B), as previously reported.^20, 21^ Muscle degeneration remained mild in 9- and 12-month-old *Mbnl1^ΔE3/ΔE3^* and *HSA^LR^* mice, respectively (*Figure* 1A and 1B). Alterations, such as internalized nuclei, vacuoles and degenerated fibres, were present in *Mbnl1^ΔE3/ΔE3^* muscle and in *gastrocnemius* muscle from *HSA^LR^* mice. In contrast, alterations were rare in *tibialis anterior* (TA) and *extensor digitorum longus* (EDL) muscles from *HSA^LR^* mice (*Figure* 1A and 1B). Both *Mbnl1^ΔE3/ΔE3^* and *HSA^LR^* muscles displayed a myotonic phenotype, as shown by the increased late relaxation time of EDL muscle after *ex vivo* stimulation (*Figure* 1C and 1D). In contrast, specific tetanic muscle force (sP0) was not affected in 3-month-old mutant mice (*Figure* 1E), consistent with previous reports showing late onset of muscle weakness.^20, 21^

**Figure 1:**
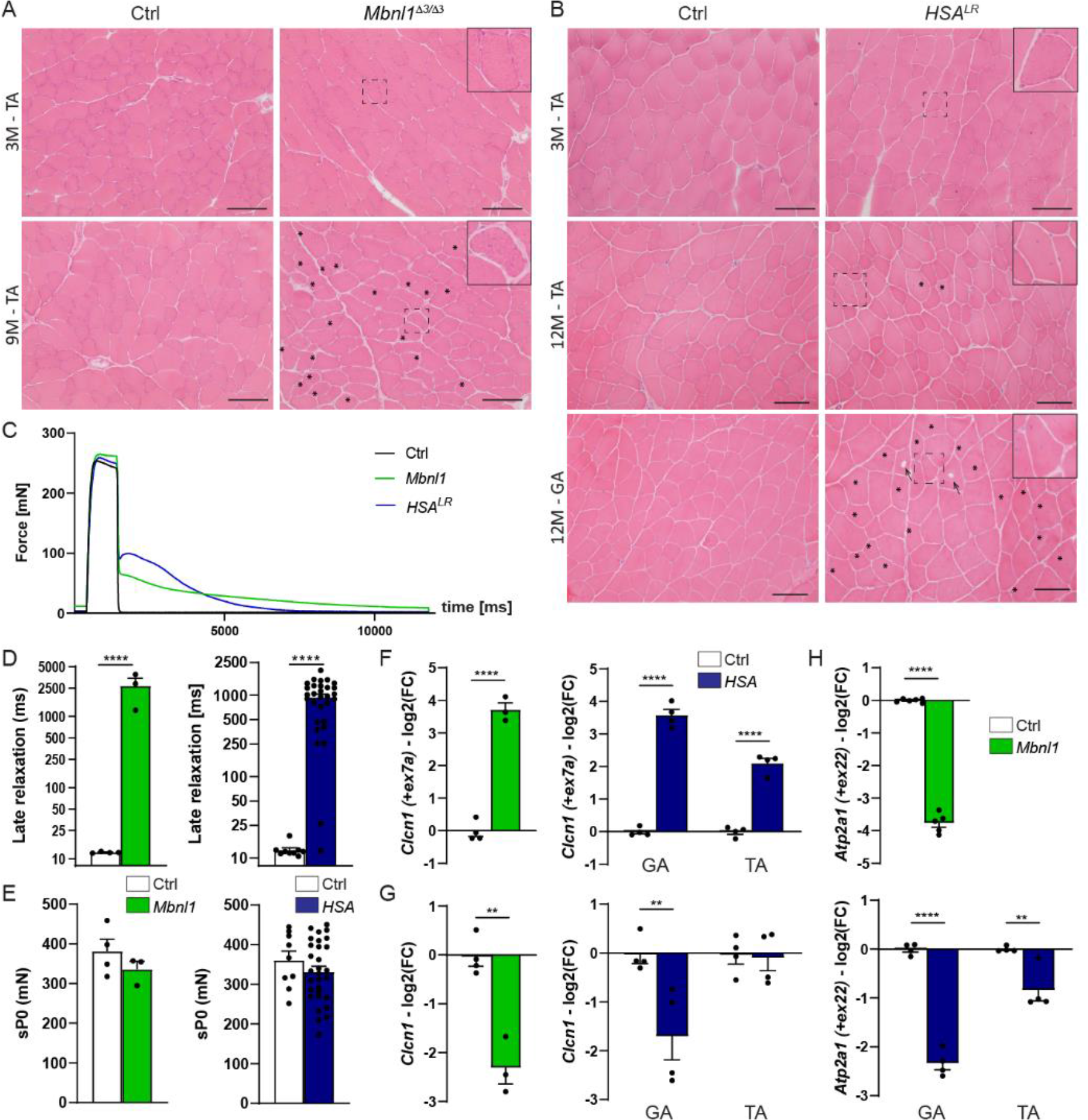
Muscle affection in *Mbnl1^ΔE3/ΔE3^* and *HSA^LR^* mice. A, B) H&E coloration reveals limited histopathologic alterations in muscles from 3- and 9-month-old *Mbnl1^ΔE3/ΔE3^* (A) and 3- and 12-month-old *HSA^LR^* mice (B). Asterisks and arrows point to internalized nuclei and vacuoles, respectively. Scale bar, 100 µm. C, D) Late relaxation time upon stimulation is increased in EDL muscle from 3-month-old *Mbnl1^ΔE3/ΔE3^* and *HSA^LR^* mice, as compared to control mice. n = 4 Ctrl / 3 *Mbnl1^ΔE3/ΔE3^*; 9 Ctrl / 30 *HSA^LR^*. E) Specific tetanic force (sP0) of EDL muscle is unchanged in 3-month-old *Mbnl1*^*ΔE3/ΔE3*^ (n = 4 Ctrl / 3 KO) and *HSA*^*LR*^ (n= 9 Ctrl / 30 *HSA*^*LR*^) mice. F-H) Quantitative RT-PCR analysis of the inclusion of *Clcn1* exon7a (F), of total *Clcn1* transcript (G), and of the inclusion of *Atp2a1* exon22 (H) in TA muscle from *Mbnl1*^*ΔE3/ΔE3*^ mice, and gastrocnemius or TA muscle from *HSA*^*LR*^ mice. Expression of spliced variants is normalised on total expression. Total expression is normalized on *Tbp* expression. n = 4 Ctrl / 3 *Mbnl1*^*ΔE3/ΔE3*^ (for Clcn1); 6 Ctrl / 5 *Mbnl1*^*ΔE3/ΔE3*^ (for Atp2a1); 4 Ctrl / 4 *HSA*^*LR*^. In all panels, data represent mean ± SEM. *p<0.05, **p<0.01, ***p<0.001, ****p<0.0001, unpaired two-tailed Student’s t test.

To evaluate DM1-associated mis-splicing, we next quantified the inclusion of exons 7a and 22 of the *Clcn1* and *Atp2a1* genes, encoding ClC-1 channel and SERCA1 (*Sarco/Endoplasmic Reticulum Ca^2+^-ATPase*), respectively. Inclusion of *Clcn1* exon 7a was strongly increased in *Mbnl1^ΔE3/ΔE3^* muscle and was accompanied by major reduction in total transcript levels of *Clcn1* (*Figure* 1F). *Clcn1* mis-splicing and down-regulation were comparable in *gastrocnemius* muscle from *HSA^LR^* mice (*Figure* 1F). In contrast, TA muscle from *HSA^LR^* mice showed milder changes in *Clcn1* splicing and no reduction in total *Clcn1* transcript levels (*Figure* 1F). Similarly, inclusion of *Atp2a1* exon 22 was abrogated in *Mbnl1^ΔE3/ΔE3^* muscle and in *gastrocnemius* muscle from *HSA^LR^* mice, while it was only reduced by half in *HSA^LR^* TA muscle (*Figure* 1F). These results confirm that both mouse models display mild muscle alterations, with similar DM1-associated phenotype observed in *Mbnl1^ΔE3/ΔE3^* muscle and *HSA^LR^ gastrocnemius* muscles, and milder changes detected in TA/EDL muscles from *HSA^LR^* mice.

### DM1 muscle shows major deregulation of CaMKIIs

Previous reports pointed to mis-splicing of *CAMK2* genes in DM1 tissues.^3–5^ However, it remains unclear which CaMKII isoforms are affected in DM1 muscle, and what are the consequences of their deregulation. CaMKIIs are encoded by the four genes *CAMK2A, 2B, 2G* and *2D*, and expressed as different splice variants. CaMKIIβ/γ/δ isoforms are expressed in skeletal muscle. Protein levels of the muscle-specific variant of CaMKIIβ (*i.e.,* CaMKIIβM) were strongly reduced in both *Mbnl1^ΔE3/ΔE3^* (*Figure* 2A) and *HSA^LR^* (*Figure* 2B) muscles. Auto-phosphorylation of CaMKIIβM (Phospho-Ser286) was decreased as well, suggesting reduced CaMKIIβM activity (*Figure* 2C). Of note, additional bands around the size of the ubiquitous isoforms of CaMKIIβ and of CaMKIIγ/δ point to the expression of alternative CaMKII isoforms in mutant muscle (*Figure* 2A and 2B).

**Figure 2:**
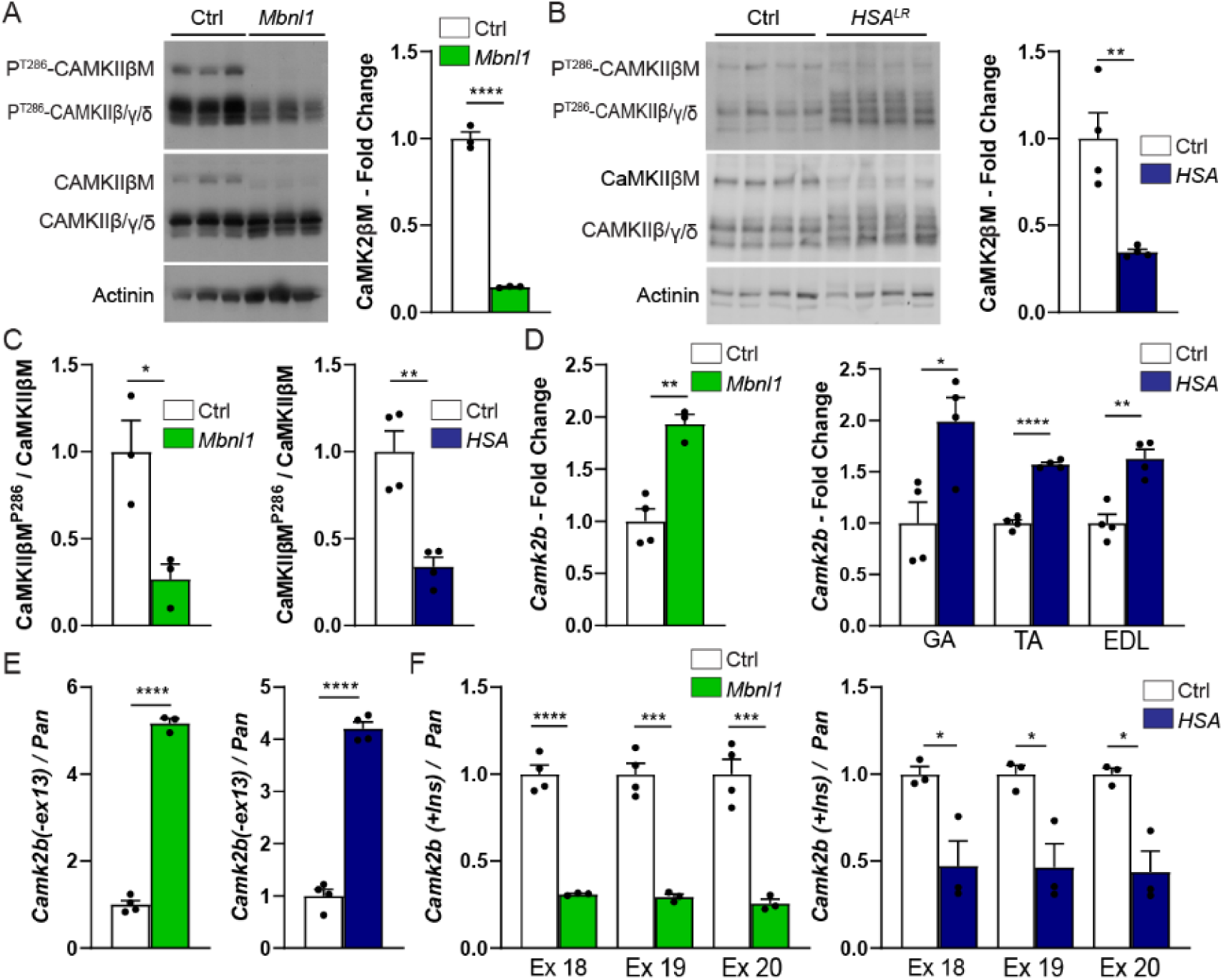
CaMKIIβ deregulation in *Mbnl1^ΔE3/ΔE3^* and *HSA^LR^* muscles. A-C) Western blot analysis of CaMKII isoforms and quantification of CaMKIIβM levels in TA muscle from 3-month-old *Mbnl1^ΔE3/ΔE3^* mice (A) and in *gastrocnemius* muscle from 3-month-old *HSA^LR^* mice (B). Quantification of CaMKIIβM phosphorylated form in *Mbnl1^ΔE3/ΔE3^* and *HSA^LR^* muscles is shown in C. Normalization is made on α-actinin (A, B) or on total CaMKIIβM (C), relative to control. n = 3 (Ctrl/*Mbnl1^ΔE3/ΔE3^*) and 4 (Ctrl/*HSA^LR^*) per group. D-F) Quantitative RT-PCR analysis of total levels of *Camk2b* (D), of the exclusion of exon 13 (E), and of the inclusion of exons 18-20 (F) in TA muscle from *Mbnl1^ΔE3/ΔE3^* mice and in *gastrocnemius*, TA and EDL muscles from *HSA^LR^* mice. Data are normalized on levels of *Tbp* (D) or on total *Camk2b* transcripts (E, F). n = 4 Ctrl / 3 *Mbnl1^ΔE3/ΔE3^*; 4 Ctrl / 4 *HSA^LR^* (D, E); 3 Ctrl / 3 *HSA^LR^* (F). All data are mean ± SEM; * p<0.05; ** p<0.01; **** p<0.0001; two-tailed unpaired Student’s t-test.

To characterize splicing events in *Camk2b, 2g* and *2d* genes in DM1 muscle, we took advantage of RNA-seq results that we obtained in *HSA^LR^ gastrocnemius* muscle. We detected significant changes in *Camk2b* transcript, with an increased exclusion of the exons 13 and 18 to 20 in *HSA^LR^* muscle, compared to control (*Figure* S1A and S1B). We confirmed the expression of *Camk2b* transcripts lacking exon 13 in *HSA^LR^* muscle by RT-PCR (*Figure* S1C and S1D). Moreover, the amplicon including exons 18 to 20, encoding the variable inserts A, B and C of CaMKIIβM, was barely detected in *HSA^LR^* muscle (*Figure* S1C and S1D). By quantitative RT-PCR, total levels of *Camk2b* transcripts were increased in both *Mbnl1^ΔE3/ΔE3^* and *HSA^LR^* muscles (*Figure* 2D). Using primers spanning *Camk2b* exons 12-14, we confirmed that expression of *Camk2b* transcripts without exon 13 was more than four times higher in mutant muscles than in controls (*Figure* 2E). Inversely, levels of *Camk2b* transcripts with exons 18 to 20 were reduced by half in *Mbnl1^ΔE3/ΔE3^* TA muscle and in *gastrocnemius* muscle from *HSA^LR^* mice (*Figure* 2F) compared to controls. Of note, TA and EDL muscles from *HSA^LR^* mice showed similar extent of *Camk2b* mis-splicing compared to *gastrocnemius* muscle (*Figure* S1E and S1F). Strikingly, the mis-splicing of exon 13 and exons 18-20 is also detected in RNA-seq data of TA muscle from DM1 patients (*Figure* S2).^28^

Transcripts encoding CaMKIIγ were also mis-spliced in *HSA^LR^* muscle. These included increased inclusion of exons 13, 15 and 19 in *Camk2g* transcripts in *HSA^LR^* muscle, as seen in RNA-seq reads (*Figure* S3A and S3B) and by RT-PCR (*Figure* S3C). Increased inclusion of *CAMK2G* exon 19 was also observed in TA biopsies from DM1 patients (*Figure* S3D).^28^ In contrast, there was no major splicing change detected for *CAMK2D* in DM1 muscle (*Figure* S4).^28^ Together, these results show that some *Camk2* transcripts are mis-spliced in DM1 muscle and that the expression pattern of CaMKIIs is altered, with a loss of the muscle-specific variant CaMKIIβM.

### Endplate fragmentation is not caused by abnormal AChR dynamics in *Mbnl1^ΔE3/ΔE3^* and *HSA^LR^* mice

As CaMKIIs are key sensors of neural activity involved in NMJ maintenance, we next analysed NMJ structures in EDL, TA and *gastrocnemius* muscles from 3-month-old and 9- or 12-month-old *Mbnl1^ΔE3/ΔE3^* and *HSA^LR^* mice. Pre- and post-synaptic compartments were stained with antibodies against Neurofilament/Synaptophysin and with α-bungarotoxin (Btx), which binds specifically to AChRs, respectively. The overall organization of the NMJs was preserved in mutant mice (*Figure* 3A and 3B). In particular, denervated endplate and abnormal axonal termination were not observed in mutant muscle. However, fragmentation of the endplate was increased in muscles from both 3- and 9-month-old *Mbnl1^ΔE3/ΔE3^* mice, compared to age-matched controls (*Figure* 3C). A similar increase in endplate fragmentation was observed in both *gastrocnemius* and TA/EDL muscles from *HSA^LR^* mice (*Figure* 3D). As the *HSA^LR^* transgene is specifically expressed in skeletal muscle, these results indicate that post-synaptic perturbations contribute to endplate fragmentation in *HSA^LR^* mice. Moreover, these NMJ alterations were detected as soon as 3 months of age, *i.e.* before changes in muscle histology, suggesting that they are primary deficit in DM1 mice and not a consequence of muscle degeneration/regeneration.

**Figure 3:**
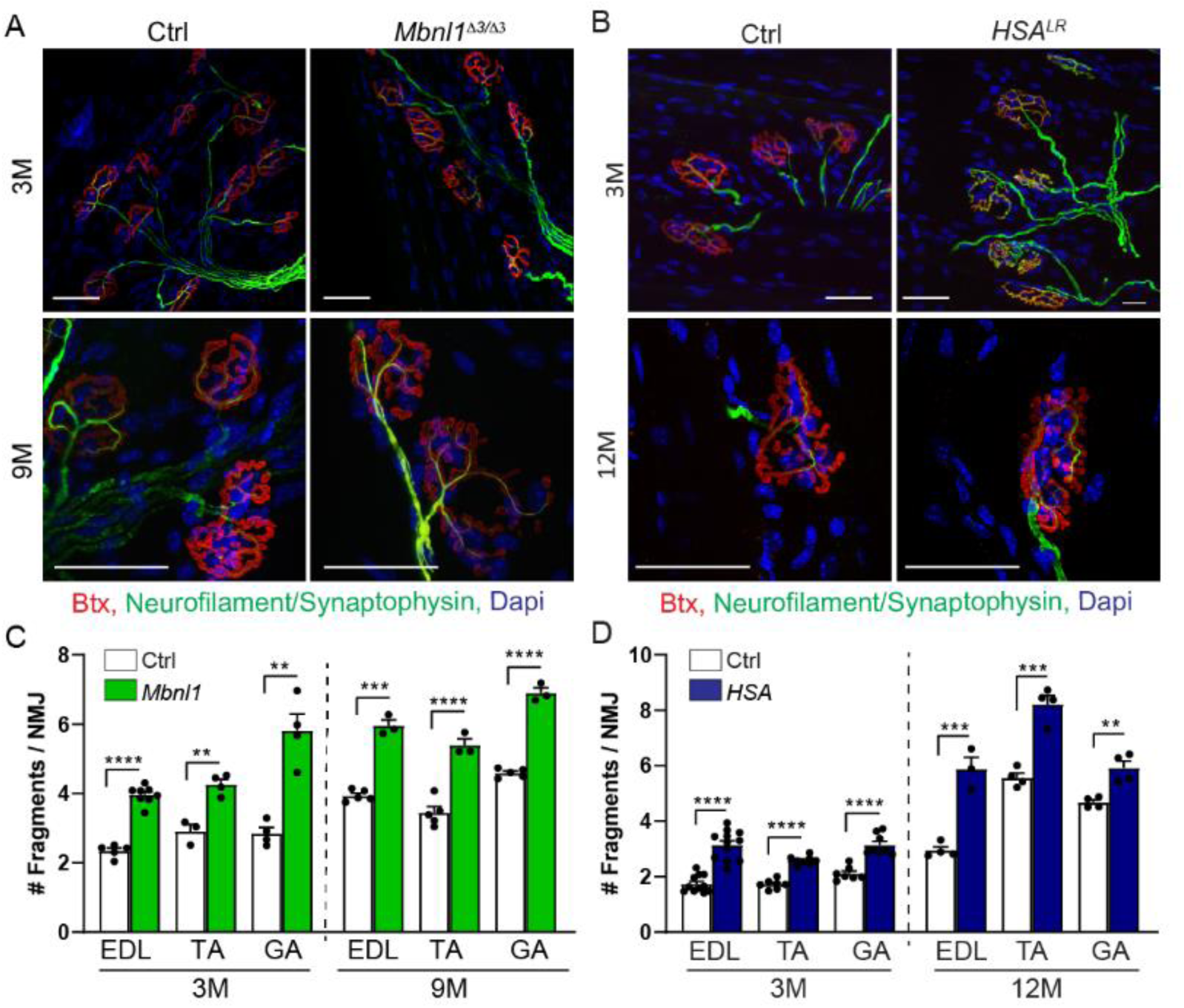
Altered NMJ maintenance in *Mbnl1^ΔE3/ΔE3^* and *HSA^LR^* mice. A, B) Fluorescent images of NMJ regions stained with α-bungarotoxin (Btx, red), antibodies against Neurofilament/Synaptophysin (green), and Dapi (blue) in EDL muscles from 3- and 9-month-old *Mbnl1^ΔE3/ΔE3^* mice (A) and 3-and 12-month-old *HSA^LR^* mice (B). Scale bar, 50 µm. C, D) Quantification of the number of fragments per NMJ in EDL, TA and *gastrocnemius* muscles from 3- and 9/12-month-old *Mbnl1^ΔE3/ΔE3^* (C) and *HSA^LR^* (D) mice. n=5/8 (EDL 3M), 3/4 (TA 3M), 5/3 (all muscles 9M) Ctrl/ *Mbnl1^ΔE3/ΔE3^* (C); 11/12 (EDL 3M), 7/8 (TA 3M), 7/8 (GA 3M), 4/3 (EDL 12M), 4/4 (TA and GA 12M) Ctrl/*HSA^LR^* (D), with more than 52 fibres per muscle. Data represent mean ± SEM. **p<0.01, ***p<0.001, ****p<0.0001, unpaired two-tailed Student’s t test.

As CaMKIIs have been shown to regulate AChR recycling at the endplate,^6^ we next evaluated AChR turnover, by labelling “old” and newly formed receptors by two sequential injections of differently labelled Btx (*Figure* 4A and 4B).^26, 29^ AChR turnover was low in all muscles, as described before for innervated conditions, with no change detected in *Mbnl1^ΔE3/ΔE3^* muscles as compared to controls (*Figure* 4C). In contrast, there was a trend towards increased AChR turnover in TA and EDL muscles from *HSA^LR^* mice (*Figure* 4D). We further determined the extent of AChR recycling in *Mbnl1^ΔE3/ΔE3^* EDL muscle, by using Btx-biotin and two sequential injections of fluorescent streptavidin (*Figure* 4B).^27^ After 1 and 3 days, the proportion of recycled AChRs was also unaltered in muscle from *Mbnl1^ΔE3/ΔE3^* mice compared to controls (*Figure* 4E and 4F). Together, these results indicate that CaMKII deregulation, and more specifically the loss of CaMKIIβM, do not alter AChR dynamics in innervated DM1 muscle.

**Figure 4:**
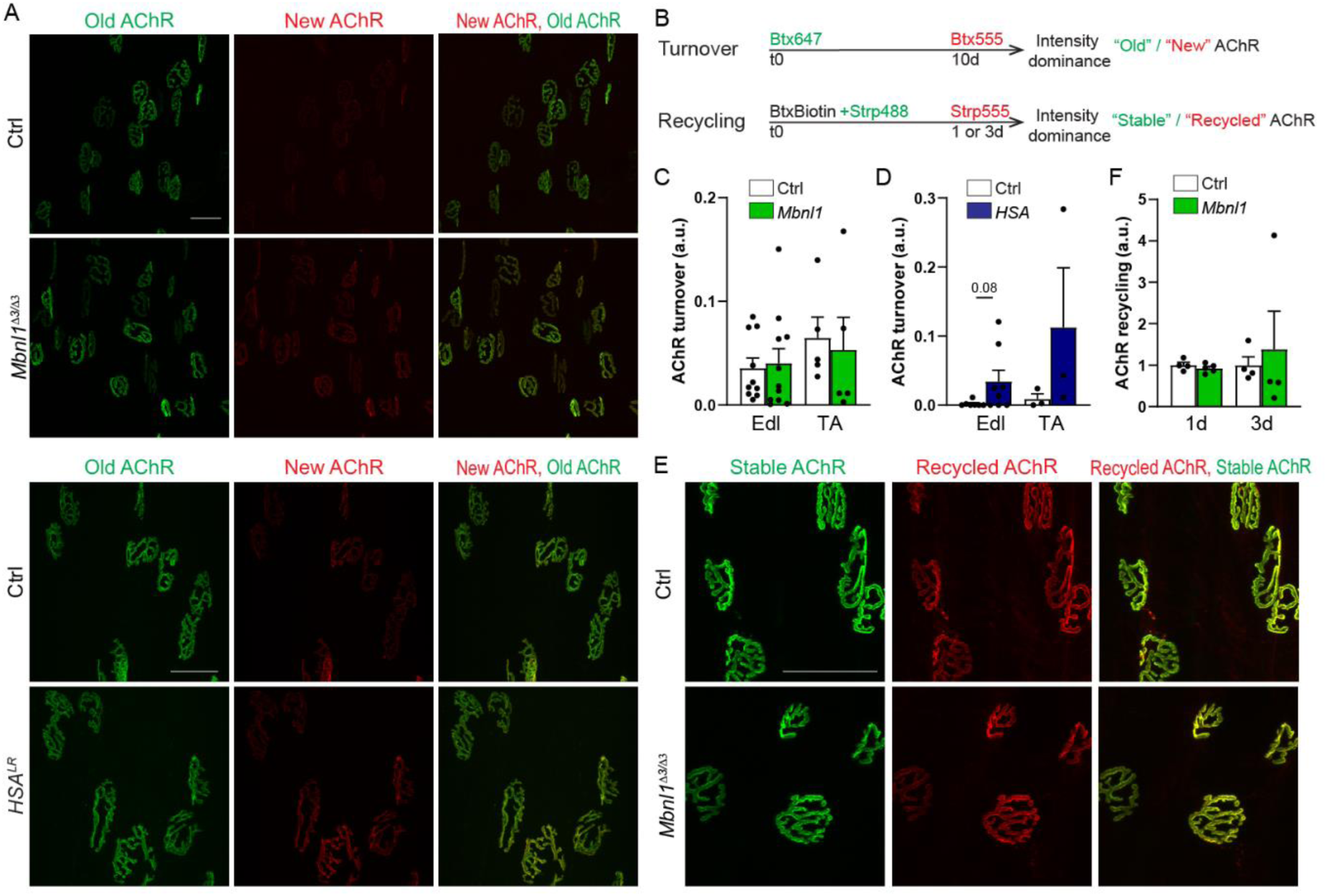
AChR dynamics is not altered in *Mbnl1^ΔE3/ΔE3^* and *HSA^LR^* muscles. A) Turnover assay in EDL and TA muscles from *Mbnl1^ΔE3/ΔE3^* and *HSA^LR^* mice. Fluorescent images show “old” (green) and “new” (red) AChRs in *Mbnl1^ΔE3/ΔE3^* and *HSA^LR^* muscles. Scale bar, 50 µm. B) Timeline of injections of α-bungarotoxin (Btx) for AChR turnover and recycling assays. *d, days; Strp, streptavidin.* C, D) AChR turnover in EDL and TA muscles from *Mbnl1^ΔE3/ΔE3^* (C) and *HSA^LR^* (D) mice. n = 10/11 (EDL) and 5/5 (TA) Ctrl/*Mbnl1^ΔE3/ΔE3^* (C); 7/8 (EDL) and 3/3 (TA) Ctrl/*HSA^LR^* (D), with more than 22 fibres per muscle. E, F) AChR recycling in EDL muscle from *Mbnl1^ΔE3/ΔE3^* mice. Fluorescent images show “stable” (green) and “recycled” (red) AChRs in *Mbnl1^ΔE3/ΔE3^* muscles (E). Scale bar, 50 µm. Quantification is given in F. n = 4/5 (1d); 4/4 (3d) Ctrl/*Mbnl1^ΔE3/ΔE3^* per group (F), with more than 28 fibres per muscle. Data are mean ± SEM; *** p<0.001; two-tailed unpaired Student’s t-test.

### HDAC4 accumulates in DM1 muscle

Another function of CaMKIIs in skeletal muscle is to repress synaptic gene expression, by inhibiting the nuclear import, and thereby the activity of HDACs, such as HDAC4.^30^ We therefore assessed the expression pattern of HDAC4 in DM1 muscle. Transcript levels of *Hdac4* were unchanged in DM1 mice, as compared to controls (*Figure* 5A). In contrast, protein levels of HDAC4 were strongly increased in both *Mbnl1^ΔE3/ΔE3^* and *HSA^LR^* muscles, as compared to controls (*Figure* 5B and 5C). While CaMKII-dependent phosphorylation of HDAC4 (Ser632) remained unchanged in *Mbnl1^ΔE3/ΔE3^* muscle (*Figure* 5B), it tended to decrease in *HSA^LR^* muscle, as compared to controls (*Figure* 5C). Consistent with a reduced inhibition of HDAC4 by CaMKIIs, HDAC4 levels were increased in the nuclear fraction of *Mbnl1^ΔE3/ΔE3^* and *HSA^LR^ gastrocnemius* muscles (*Figure* 5D). Of note, HDAC4 accumulation remained barely detectable in mutant muscle by immunostaining (*Figure* 5E) and it was not sufficient to increase total levels of acetylated histones, as reported upon denervation (*Figure* S5A and S5B). Consistent with higher HDAC4 activity, transcript levels of *Dach2*, a gene directly repressed by HDAC4 (*Figure* S5C), tended to decrease in DM1 muscles (*Figure* 5F). In contrast, expression of *Mitr* was not repressed in mutant muscles, and transcript levels of *Myog,* encoding myogenin, remained unchanged in *Mbnl1^ΔE3/ΔE3^* and *HSA^LR^* muscles, as compared to controls (*Figures* 5F, S5C and S5D). Hence, CaMKII deregulation may trigger accumulation of HDAC4 and a moderate increase in its activity in DM1 muscle.

**Figure 5:**
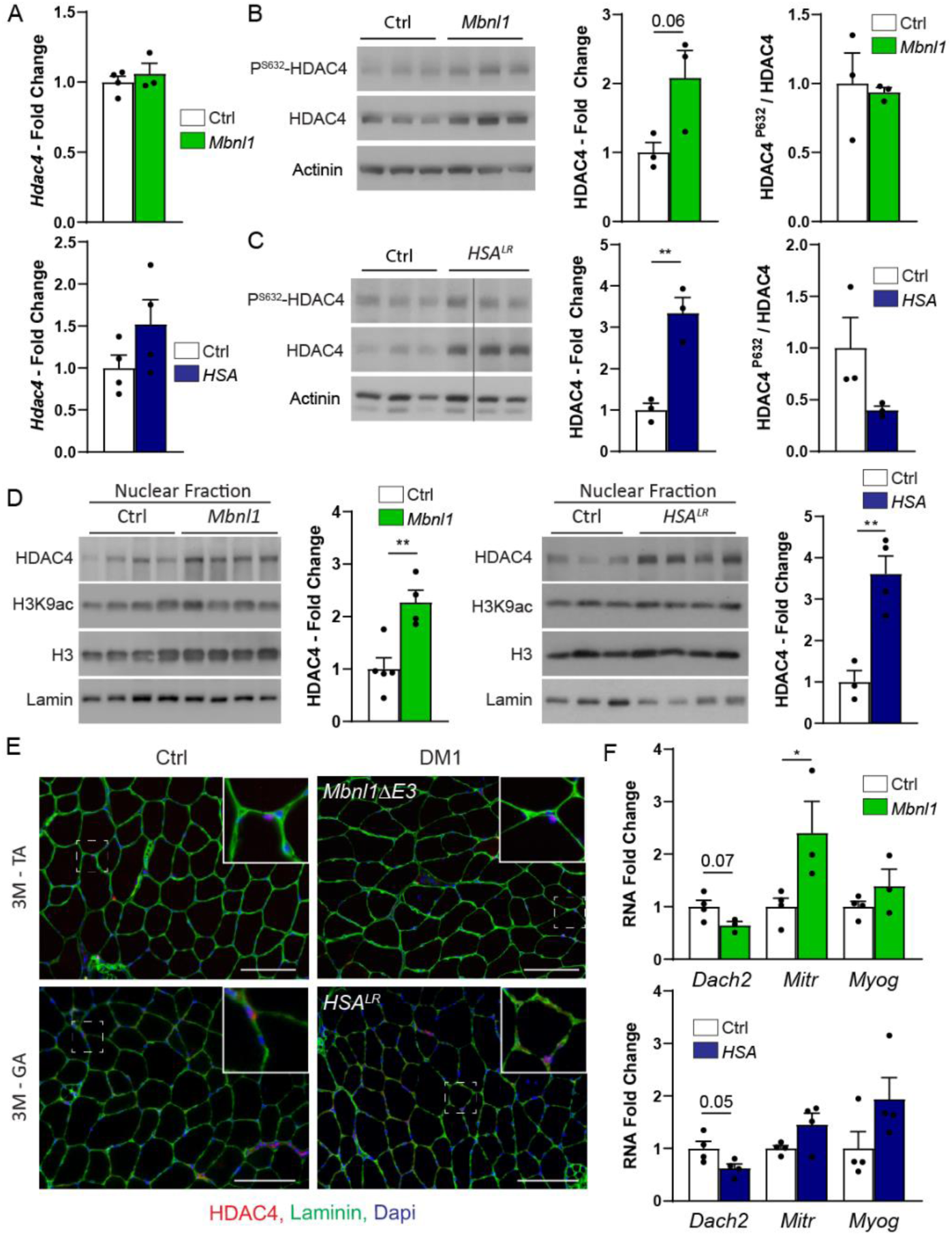
Changes in HDAC4 signalling pathway in *Mbnl1^ΔE3/ΔE3^* and *HSA^LR^* muscles. A) Quantitative RT-PCR analysis of *Hdac4* in *Mbnl1^ΔE3/ΔE3^* and *HSA^LR^* muscles. n = 4/3 Ctrl/*Mbnl1^ΔE3/ΔE3^* and 4/4 Ctrl/ *HSA^LR^*. B, C) Western blot analysis of HDAC4 and its phosphorylated form (Ser632) in total protein lysate of *Mbnl1^ΔE3/ΔE3^* (B) and *HSA^LR^* (C) muscles. Quantification of total and phosphorylated levels is given for *Mbnl1^ΔE3/ΔE3^* (B) and *HSA^LR^* (C) muscles. Total and phosphorylated levels are normalized to α-actinin and total HDAC4, respectively, and relative to control. n = 3 per group. D) Western blot analysis of HDAC4 and acetylated histones (H3K9) in nuclear fraction of *gastrocnemius* muscle from *Mbnl1^ΔE3/ΔE3^* mice and *HSA^LR^* mice. Quantifications of HDAC4 and H3K9 levels in *Mbnl1^ΔE3/ΔE3^* and *HSA^LR^* muscles are given in (D) and in Figure S5, respectively. Levels are normalized to lamin, and relative to control. n = 5/4 Ctrl/*Mbnl1^ΔE3/ΔE3^* and 3/4 Ctrl/*HSA^LR^*. E) Fluorescent image of *Mbnl1^ΔE3/ΔE3^* and *HSA^LR^* muscles stained with antibodies against HDAC4 (red), Laminin (green) and Dapi (blue). Scale bar, 100 µm. Higher magnification panel shows positive myonuclei. F) Quantitative RT-PCR analysis of activity-dependent genes, *Dach2*, *Mitr* and *Myog* in TA muscle from *Mbnl1^ΔE3/ΔE3^* mice and in *gastrocnemius* muscle from *HSA^LR^* mice. Transcript levels are normalized to *Tbp* and relative to control. n = 4/3 Ctrl/*Mbnl1^ΔE3/ΔE3^* and 4/4 Ctrl/*HSA^LR^*. All data are mean ± SEM; * p<0.05; ** p<0.01; *** p<0.001; two-tailed unpaired Student’s t-test.

### Expression of synaptic genes and fibre type composition are altered in DM1 muscle

We next assessed the consequences of CaMKII/HDAC4 deregulation in DM1 muscle on known activity-dependent signalling pathways (*Figure* S5C). We first measured the transcriptional expression of *Musk*, *Chrna1*, *Chrne* and *Chrng* genes, encoding MUSK and the α, ε and γ subunits of AChR, respectively. During muscle development, expression of *Musk* and *Chrna1* becomes restricted to sub-synaptic myonuclei upon muscle innervation.^31^ Simultaneously, *Chrng* transcripts are downregulated, while *Chrne* starts to be expressed in sub-synaptic nuclei (AChRε subunits replace AChRγ subunits). In non-synaptic regions of muscle fibres, synaptic gene repression is mediated by CaMKII, and dependent on HDAC4/5 inhibition (*Figure* S5C).^8, 32^ This adult gene expression pattern depends on neural activity, as denervation reverts it back to a developmental pattern. As shown in Figure 6A, transcript levels of *Musk, Chrna1, Chrne* and *Chrng,* were increased in *Mbnl1^ΔE3/ΔE3^* muscle and in *gastrocnemius* from *HSA^LR^* mice. Of note, their levels were less or not changed in EDL and TA muscles from *HSA^LR^* mice, as compared to controls (*Figure* S6A and S6B).

**Figure 6:**
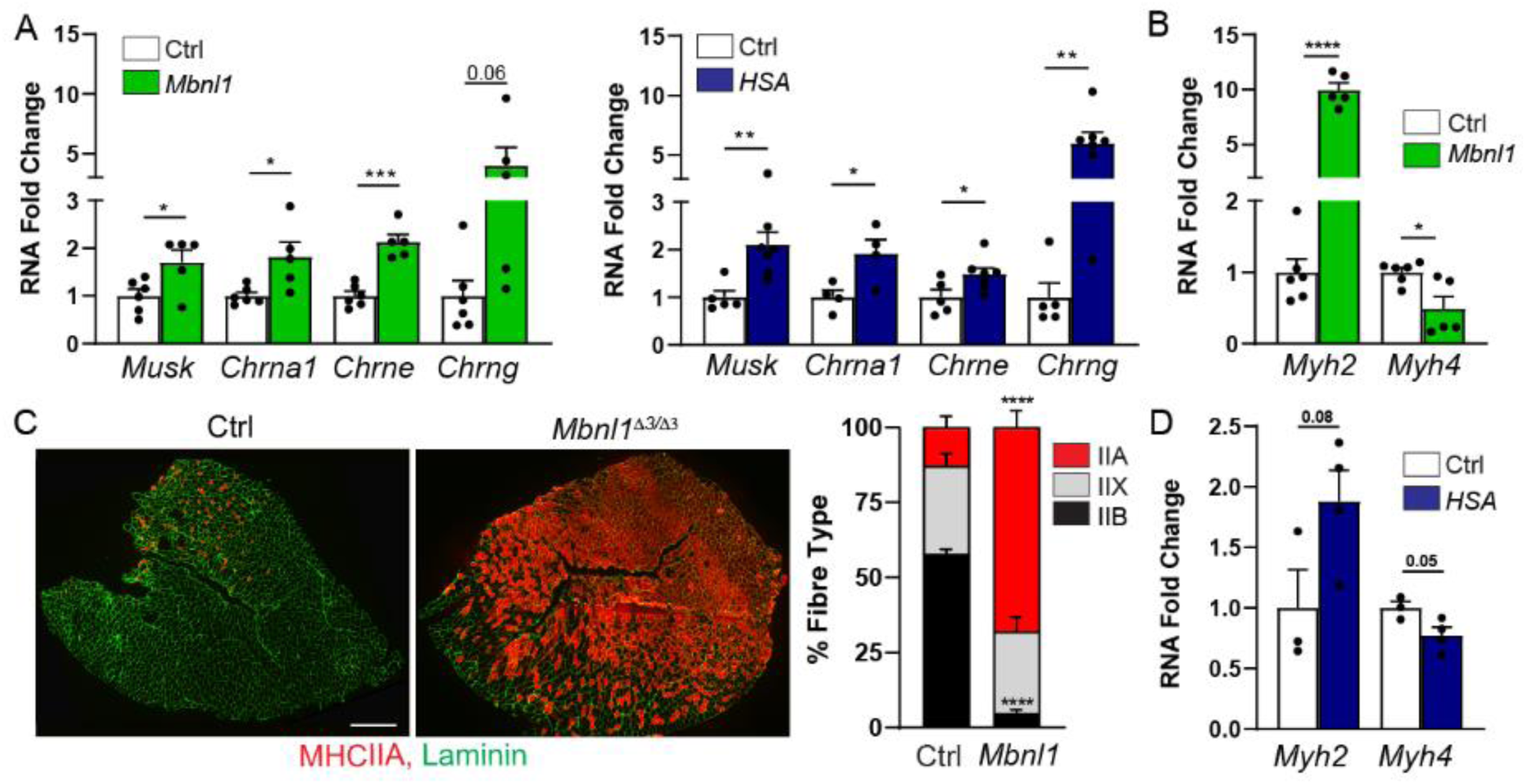
DM1 mice display deregulation of activity-dependent signalling pathways. A) Quantitative RT-PCR analysis of *Musk, Chrna1, Chrne* and *Chrng* in TA muscle from 3-month-old *Mbnl1^ΔE3/ΔE3^* mice and in *gastrocnemius* muscle from 3-month-old *HSA^LR^* mice. Levels are normalized to *Tbp*. n=6/5 Ctrl/ *Mbnl1^ΔE3/ΔE3^* and 5/7 Ctrl/*HSA^LR^* (except for *Chrna1*, n=4). B) Quantitative RT-PCR analysis of *Myh2* and *Myh4*, encoding MHC2A and MHC2B, in *Mbnl1^ΔE3/ΔE3^* muscle. Transcript levels are normalized to *Tbp* and relative to control. n=6/5 Ctrl/*Mbnl1^ΔE3/ΔE3^*. C) Fluorescent images of control and *Mbnl1^ΔE3/ΔE3^* muscles, stained with antibodies against MHC2A (red) and Laminin (green), and quantification of the proportion of type IIA, IIX and IIB fibres in control and mutant muscles. Scale bar, 500 µm. D) Quantitative RT-PCR analysis of *Myh2* and *Myh4* in *HSA^LR^* TA muscle. Transcript levels are normalized to *Tbp* and relative to control. n=3/4 Ctrl/ *HSA^LR^*. All data are mean ± SEM; * p<0.05; ** p<0.01; *** p<0.001; two-tailed unpaired Student’s t-test.

HDAC4 is also known to regulate the transcription of *Myh* genes, encoding myosin heavy chains (MHC), and thereby to promote a switch to type IIA fibres in TA muscle upon denervation.^8^ Expression of *Myh2*, encoding MHCIIA, was strongly increased in *Mbnl1^ΔE3/ΔE3^* innervated muscle, as compared to control (*Figure* 6B). In contrast, expression of *Myh4*, encoding MHCIIB, was reduced in DM1 muscle (*Figure* 6B). Consistently, innervated TA muscle from *Mbnl1^ΔE3/ΔE3^* mice displayed major accumulation of type IIA fibres and a loss of type IIB fibres compared to controls (*Figure* 6C). Similarly, we have previously unveiled a mild switch towards type IIA fibres in TA muscle from *HSA^LR^* mice.^23^ There was also a tendency towards increased transcript levels of *Myh2* and reduced expression of *Myh4* in *HSA^LR^* TA muscle, as compared to controls (*Figure* 6D). Thus, CaMKII-dependent deregulation of HDAC4 may perturb activity-dependent signalling pathways, with increased synaptic gene expression and a switch towards slower fibres in DM1 muscle.

### *Mbnl1^ΔE3/ΔE3^* muscle responds to denervation

To get further insights on the deregulation of activity-dependent signalling pathways in DM1 muscles, we assessed their response to neural inactivity. To this end, we challenged 3-month-old *Mbnl1^ΔE3/ΔE3^* and *HSA^LR^* mice by cutting the sciatic nerve to obtain a complete denervation of hind limb muscles. Unexpectedly, we observed that *HSA^LR^* mice lose the expression of the *HSA* transgene after 3 days of denervation (*Figure* S7A). Consequently, accumulation of ribonuclear foci and mis-splicing of *Clcn1* and *Camk2b* were reduced in denervated *HSA^LR^* muscle, as compared to innervated muscle (*Figure* S7B-D). Because of this, we limited the analysis to the *Mbnl1^ΔE3/ΔE3^* mouse line.

After 3 days of denervation, we observed an efficient increase in transcript and protein levels of HDAC4 in denervated muscle from both mutant and control mice (*Figure* 7A). Consistently, there was a major accumulation of HDAC4 in nuclei of both DM1 and control denervated muscles (*Figure* 7B). Of note, the loss of CaMKIIβM persisted upon denervation in *Mbnl1^ΔE3/ΔE3^* muscle (*Figure* 7A). Expression of the direct targets of HDAC4, *Dach2* and *Mitr*, was similarly repressed in *Mbnl1^ΔE3/ΔE3^* and control denervated muscles (*Figure* 7C). Moreover, expression of the atrogenes *Fbxo32* and *Trim63*, which are indirectly induced by HDAC4 at this stage,^33^ also reached similar transcript levels in *Mbnl1^ΔE3/ΔE3^* and control muscles (*Figure* S8A). We and others have reported that the anabolic pathway Akt/mTORC1 (*mammalian Target Of Rapamycin Complex 1*) is deregulated in DM1 muscle,^23^ and that it is activated upon denervation.^26^ Thus, we assessed whether Akt/mTORC1 activity is perturbed in DM1 denervated muscle. At 3 days of denervation, there was no change in the levels of the phosphorylated active form of Akt (Akt^P473^) in both *Mbnl1^ΔE3/ΔE3^* and control muscles, while levels of the active form of S6 (S6^P235^) increased similarly in both denervated muscles (*Figure* S8B). In parallel, levels of the autophagic marker LC3II remained largely unchanged in *Mbnl1^ΔE3/ΔE3^* and control muscles, suggesting that autophagy is not strongly affected (*Figure* S8B). These results suggest that both the HDAC4 and mTORC1 signalling pathways are efficiently induced upon denervation in DM1 muscle.

**Figure 7:**
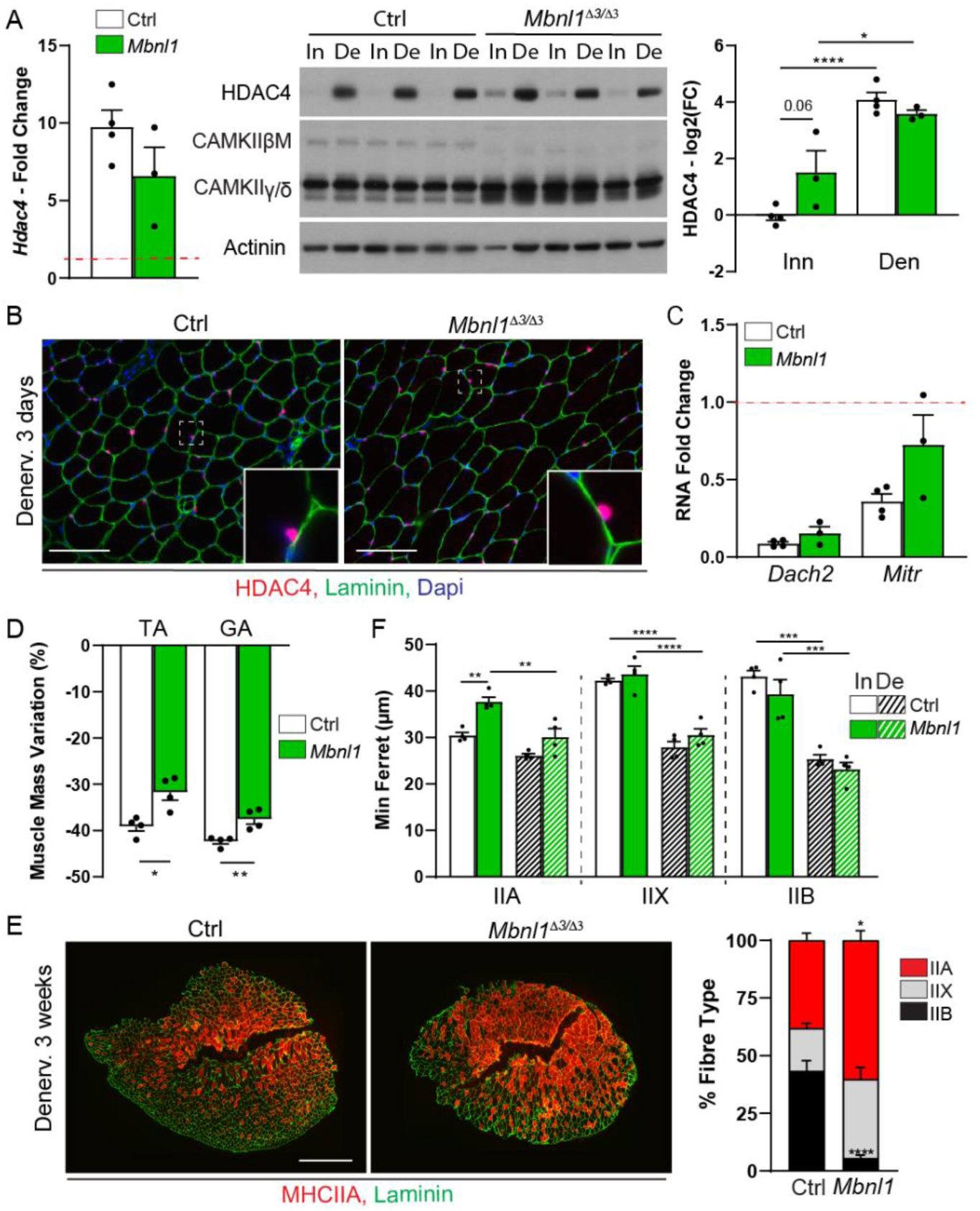
*Mbnl1^ΔE3/ΔE3^* muscle shows efficient HDAC4 induction and resistance to atrophy upon denervation. A) Quantitative RT-PCR of *Hdac4* transcript and Western blot analysis of HDAC4 protein in innervated (In) and denervated (De, 3 days) muscles from control and *Mbnl1^ΔE3/ΔE3^* mice. Transcript levels are normalized to *Tbp* and relative to control innervated. Levels in the innervated control muscle are shown by the red dotted line. Protein levels of HDAC4 are normalized to α-actinin, relative to control and expressed as the log2(Fold Change). n=4 (qPCR) and 4/3 (WB). B) Fluorescent image of *Mbnl1^ΔE3/ΔE3^* denervated muscle stained with antibodies against HDAC4 (red), Laminin (green) and Dapi (blue). Scale bar, 100 µm. Higher magnification panel shows positive myonuclei. C) Quantitative RT-PCR analysis of *Dach2* and *Mitr* in TA muscle from control and *Mbnl1^ΔE3/ΔE3^* mice 3 days post-denervation. Levels are normalized to *Tbp* and relative to control innervated. n=4 per group. Levels in the innervated control muscle are shown by the red dotted line. D) Mass variation after 3 weeks of denervation in control and *Mbnl1^ΔE3/ΔE3^* mice, for TA and *gastrocnemius* (GA) muscles. n = 4 per group. E) Fluorescent images of denervated muscles from control and *Mbnl1^ΔE3/ΔE3^* mice, stained with antibodies against type IIA myosin heavy chain (MHC2A - red) and Laminin (green) and quantification of fibre type proportion in control and mutant denervated muscles. Scale bar, 500 µm. n = 4 mice per group. F) Quantification of fibre minimum ferret in innervated and 3-week-denervated muscles from control and *Mbnl1^ΔE3/ΔE3^* mice. n = 4 mice per group. All data are mean ± SEM; * p<0.05; ** p<0.01; *** p<0.001; **** p<0.0001; two-tailed unpaired Student’s t-test (D) and two-way ANOVA with a Tukey’s post-hoc analysis (A, F).

Interestingly, after prolonged denervation, the loss of muscle mass was significantly less in *Mbnl1^ΔE3/ΔE3^* mice, as compared to control mice (*Figure* 7D). Of note, nerve injury did not aggravate muscle degeneration in *Mbnl1^ΔE3/ΔE3^* mice (*Figure* S8C). To get insights on this resistance to atrophy, we evaluated muscle fibre type and size after 3 weeks of denervation. Control TA muscle shifted to type IIA fibres upon denervation, reaching similar proportion of type IIA fibres as observed in *Mbnl1^ΔE3/ΔE3^* innervated muscle. Fibre type proportion remained unchanged in *Mbnl1^ΔE3/ΔE3^* denervated muscle (*Figure* 7E). The minimum ferret of type IIA fibres was increased in *Mbnl1^ΔE3/ΔE3^* innervated muscle as compared to control innervated muscle, but it significantly decreased upon denervation in mutant mice (*Figure* 7F and *Figure* S8D). In parallel, the size of type IIX and IIB fibres was similar in *Mbnl1^ΔE3/ΔE3^* and control innervated muscles, and it strongly decreased upon denervation in both mutant and control mice (*Figure* 7F and *Figure* S8D). Hence, the predominance of type IIA fibres in mutant mice and their relative resistance to denervation-induced atrophy as compared to type IIX/B fibres (*Figure* 7F), may explain why DM1 mice showed limited loss of muscle mass upon nerve injury.

### Endplate remodelling is impaired in *Mbnl1^ΔE3/ΔE3^* muscle upon denervation

Synaptic remodelling upon denervation includes a strong increase in AChR turnover in the sub-synaptic region, together with major up-regulation of synaptic genes throughout the fibre.^34^ Release of synaptic gene repression in non-synaptic muscle regions is thought to be dependent on the induction of HDAC4.^8, 32^ Importantly, despite the efficient induction of HDAC4 in *Mbnl1^ΔE3/ΔE3^* denervated muscle, up-regulation of *Myog*, *Musk, Chrna1* and *Chrng* was hampered in mutant muscle, as compared to controls (*Figure* 8A). Moreover, the expression of *Chrne* remained abnormally high in *Mbnl1^ΔE3/ΔE3^* muscle upon denervation (*Figure* 8B). This suggests that HDAC4-independent mechanisms likely contribute to the defective response of DM1 muscle to denervation.

**Figure 8:**
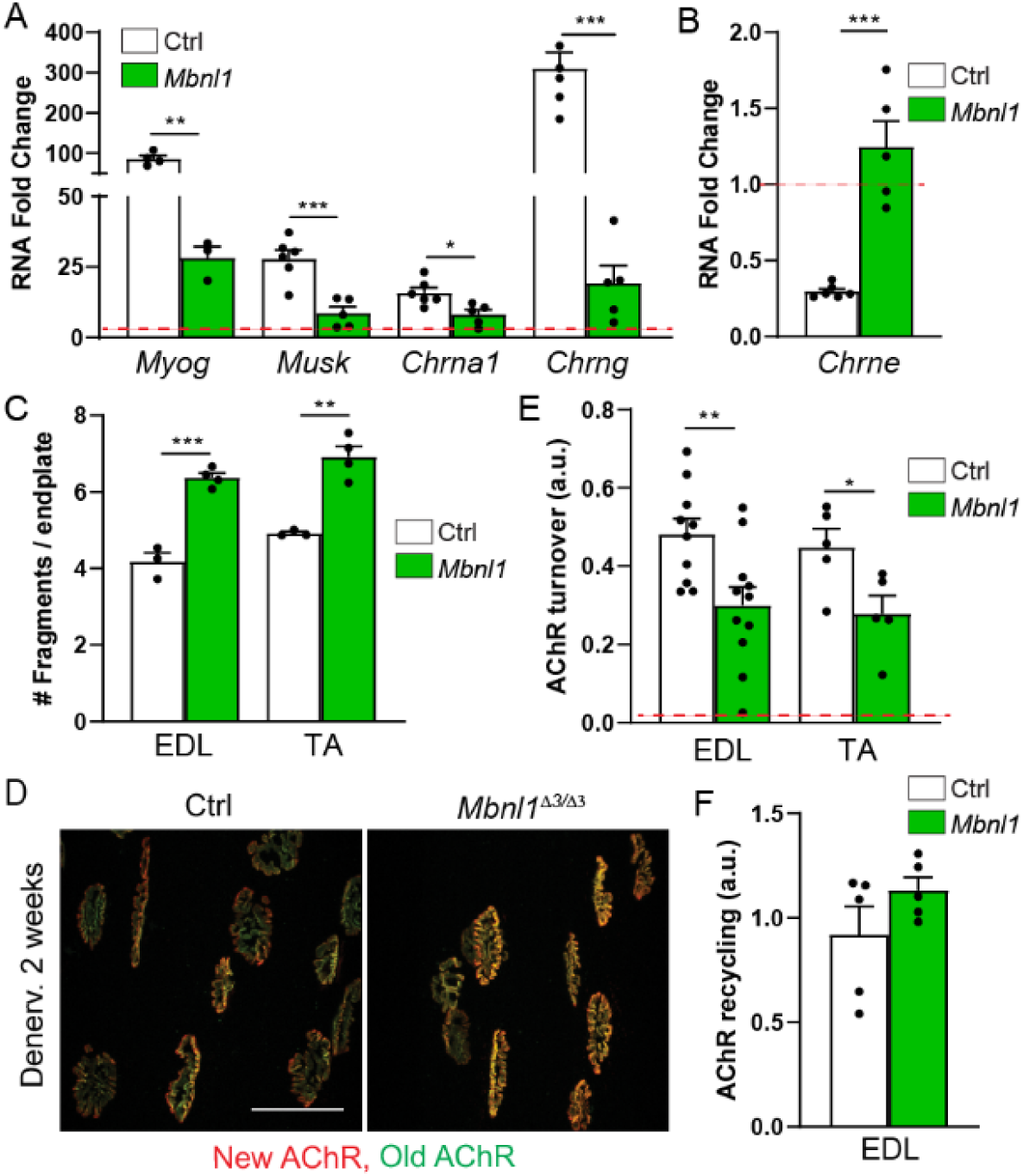
Denervation-induced changes in the expression and dynamics of synaptic proteins are impaired in *Mbnl1^ΔE3/ΔE3^* mice. A, B) Quantitative RT-PCR analysis of activity-dependent genes, *Myog, Musk, Chrna1* and *Chrng* (A), and *Chrne* (B), in TA muscle from control and *Mbnl1^ΔE3/ΔE3^* mice 3 days post-denervation. Levels are normalized to *Tbp* and relative to control innervated. n = 4 per group. Levels in the innervated control muscle are shown by the red dotted line. C) Quantification of endplate fragmentation in EDL and TA muscles from *Mbnl1^ΔE3/ΔE3^* mice after 3 weeks of denervation. n = 3/4 Ctrl/KO. D-F) AChR turnover (D, E) and recycling (F) in denervated muscle from *Mbnl1^ΔE3/ΔE3^* and control mice. Fluorescent images of “old” and “new” receptors are shown in (D). Scale bar, 50 µm. Quantification of AChR turnover and of AChR recycling is giver in (E) and (F), respectively. n = 10/11 (EDL) and 5/5 (TA) Ctrl/*Mbnl1^ΔE3/ΔE3^* (E); n=5 per group (F). Data are mean ± SEM; ** p<0.01; *** p<0.001; two-tailed unpaired Student’s t-test.

To assess the consequences of this defect, we evaluated changes at the endplate after 2 weeks of denervation. At this stage, endplate fragmentation increased and remained higher in EDL and TA muscles from *Mbnl1^ΔE3/ΔE3^* mice, compared to controls (*Figure* 8C). To go further, we quantified AChR turnover by labelling receptors 5 days after nerve injury and assessing their turnover 10 days later. In control muscle, AChR turnover increased drastically after denervation (*Figure* 8D and 8E), as previously reported.^35^ In *Mbnl1^ΔE3/ΔE3^* denervated muscle, AChR turnover was reduced as compared to control denervated muscle (*Figure* 8D and 8E). As there was no difference in AChR recycling between mutant and control denervated muscles (*Figure* 8F), reduced AChR turnover in DM1 muscle likely arose from reduced insertion and/or increased degradation of AChRs. Together, these results point to an impaired response of *Mbnl1^ΔE3/ΔE3^* mice to denervation, with reduced AChR formation, which may depend on CaMKII deficiency but unlikely on HDAC4 deregulation.

## DISCUSSION

Although mis-splicing events are at the basis of DM1 pathogenesis, it is not entirely clear how the consecutive perturbations lead to muscle alterations. NMJ deteriorations have been described in muscle biopsies from DM1 patients and in muscle from DM1 mouse models. However, it remains unknown what mechanisms underlie these perturbations and whether these defects arise from DM1-related changes in the muscle or in the nerve. Here, we showed that NMJs are affected in *Mbnl1^ΔE3/ΔE3^* and *HSA^LR^* mice, two well-characterized mouse models for DM1. We established that CaMKIIs and activity-dependent signalling pathways are disrupted in *Mbnl1^ΔE3/ΔE3^* and *HSA^LR^* muscles, which may contribute to endplate destabilization and to the abnormal response of mutant muscles to denervation.

Signs of NMJ deterioration, without denervation of muscle fibres, have been reported in muscle biopsies from DM1 patients, as well as in DMSXL and *Mbnl1/2*-deficient mice.^10–13, 16, 36^ As the motor neuron is also affected in these mouse models, it is unclear whether the defects arise from pre- or post-synaptic perturbations. Here, we found endplate fragmentation in the muscles of *Mbnl1^ΔE3/ΔE3^* and *HSA^LR^* mice at different ages. As *HSA^LR^* mice express the transgene carrying the (CTG) repeats only in muscle, it is likely that perturbations in the post-synaptic compartment, *i.e.* the muscle, are responsible for NMJ deterioration in DM1. Importantly, endplate fragmentation was similar between TA, EDL and *gastrocnemius* muscles, which are affected differentially in *HSA^LR^* mice. Especially TA and EDL muscles showed minor signs, if not none, of muscle degeneration, suggesting that endplate destabilization is causally unrelated to DM1-associated muscle degeneration. In contrast, NMJ deterioration may contribute, and eventually precede, muscle atrophy, weakness and fatigue observed in DM1 muscle.

Seeking for pathomechanisms that may compromise NMJ integrity in DM1, we examined the potential role of CaMKII deregulation. Mis-splicing in *Camk2b, 2d* and *2g* has been reported in DM1 patients, as well as in mouse models.^3–5^ In particular, abnormal exclusion of *Camk2b* exon 13 appeared as one of the most important splicing changes detected in DM1 tissues. Although pathophysiological consequences have been investigated in the brain, the consequences of CaMKII deregulation in skeletal muscle have not yet been analysed. We here report that the three exons specifically included in the muscle-specific CaMKIIβ isoform are excluded in *HSA^LR^* and *Mbnl1^ΔE3/ΔE3^* muscles, indicating that *Camk2* splicing is MBNL1-dependent. Of note, in *HSA^LR^* mice, *Camk2b* mis-splicing was detected in both *gastrocnemius* and TA/EDL muscles. Together with the exon 13 and 16, exons 18-20 are part of the highly variable region of the *Camk2b* gene, which allows the expression of tissue-specific variants. Especially, CaMKIIβM was shown to be the only isoform accumulating at the NMJ.^6^ Consistent with the abnormal splicing of *Camk2b* in DM1 muscle, CaMKIIβM was not detected in *Mbnl1^ΔE3/ΔE3^* and HSA^LR^ muscles. However, despite the known role of CaMKIIs in regulating AChR recycling, we did not find perturbation in AChR dynamics at the endplate in DM1 muscle.

CaMKIIs have also been shown to contribute to synaptic gene repression in non-synaptic regions of muscle fibres, by inhibiting HDAC4 signalling pathway and myogenin activity.^7, 32^ We unveiled that HDAC4 accumulates in myonuclei from DM1 muscle. Increased HDAC4 activity may lead to the up-regulation of synaptic genes and to the fibre type switch observed in DM1 muscle. Of note, these perturbations did not arise from spontaneous denervation of mutant muscle fibres. As there was no major change in the expression of MITR/HDAC9 and myogenin, which are thought to mediate some of the effects of HDAC4, we cannot rule out that up-regulation of activity-dependent genes in DM1 muscle involves CaMKII/HDAC4-independent mechanisms. Alternatively, changes detected in DM1 muscle may occur predominantly at the endplate, as expression of synaptic genes in the sub-synaptic region is independent from HDAC4/myogenin in innervated muscle.^31^ Synaptic gene up-regulation was detected in *Mbnl1^ΔE3/ΔE3^* muscle and in *HSA^LR^ gastrocnemius* muscle, but not in TA/EDL muscles from *HSA^LR^* mice. It is therefore unlikely that synaptic gene up-regulation causes the endplate fragmentation observed in all muscles. Of note, the fibre type switch previously reported in *HSA^LR^* muscle ^23^ was exacerbated in *Mbnl1^ΔE3/ΔE3^* muscle. This suggested that perturbations both in the muscle and in non-muscle tissues (*e.g.,* the nerve) contribute to these changes.

To get further insights on the capacity of DM1 muscle to regulate activity-dependent signalling and to maintain their endplates, we challenged *Mbnl1^ΔE3/ΔE3^* and *HSA^LR^* mice with nerve injury. Unexpectedly, transgene expression, driven by the *HSA* promoter, was lost in *HSA^LR^* muscle upon denervation, which limited the analysis. In *Mbnl1^ΔE3/ΔE3^* mice, the increase in AChR turnover triggered by denervation was reduced compared to control denervated muscle. Consistently, up-regulation of synaptic genes was hampered in mutant muscle, which may limit the incorporation of new receptors at the membrane. Up-regulation of synaptic genes upon denervation relies on the release of *Myog* expression, through the repression of *Mitr/Hdac9* and *Dach2* by HDAC4.^8, 37, 38^ Expression of these two repressors was efficiently reduced upon denervation in DM1 muscle, which was consistent with the major accumulation of HDAC4 in mutant denervated muscle. Hence, although we cannot rule out the contribution of CaMKII deregulation in the defective up-regulation of synaptic genes in DM1 denervated muscle, it is unlikely to occur *via* deregulation of HDAC4 signalling. Deregulation of distinct CaMKII isoforms may contribute to the different changes observed in DM1 muscle in innervated and denervated conditions, respectively. Changes in the activity of other repressors of *Myog*, such as Msy3/Ybx3 ^39^ or CtBP1 ^40^, may be involved as well in the defects observed in DM1 muscle.

In conclusion, our study identified NMJ deterioration as an integral part of muscle dysfunction in DM1, which likely involves muscle perturbations in activity-dependent pathways. Especially, deregulation of CaMKII isoforms may be a key event in endplate destabilization, as well as in DM1 pathogenesis in muscle and non-muscle tissues.

## METHODS

### Mice

Homozygous mice of the mouse line LR20b carrying about 250 (CTG) repeats within the *HSA* transgene (*HSA^LR^*) were obtained from Thornton and colleagues (University of Rochester Medical Centre, Rochester, New York, USA^).20^ Mice of the corresponding background strain (FVB/N) were used as control. Mice were genotyped for *HSA^LR^* transgene by quantification of *ACTA1* levels normalized to endogenous actin (mouse *Acta1*) in genomic DNA. Mice from the *Mbnl1ΔE3* line were obtained from Swanson and colleagues (College of Medicine, University of Florida, Gainesville, Florida, USA).^21^ *Mbnl1^+/+^* littermates were used as control. Mice from the *Mbnl1ΔE3* line were genotyped for exon 3 depletion in the *MBNL1* locus. Mice were maintained in a conventional specific-pathogen-free facility with a fixed light cycle (23°C, 12-hour dark-light cycle). All animal studies were performed in accordance with the European Union guidelines for animal care and approved by the Veterinary Office of the Cantons of Basel city (application number 2601) and Geneva (application number GE220).

### Muscle force and relaxation

*In vitro* force measurement of EDL muscle and late relaxation time evaluation were conducted as previously described.^22, 23^

### Western blotting

Muscles powdered in liquid nitrogen were lysed in cold RIPA+ buffer (50 mM Tris-HCl pH 8, 150 mM NaCl, 1% NP-40, 0.5% sodium deoxycholate, 0.1% SDS, 1% Triton X, 10% glycerol, phosphatase and protease inhibitors). Subcellular fractionation was done according to Dimauro et al. (2012).^24^ Following dosage (BCA Protein Assay, Pierce), proteins were separated on SDS-polyacrylamide gels and transferred to nitrocellulose membrane. Blots were blocked in TBS, 3% BSA, 0.1% Tween-20, and incubated overnight at 4°C with primary antibodies, then for 2 hours with HRP-labelled secondary antibodies. Immunoreactivity was detected using the ECL Western blot detection reagent LumiGLO (KPL) and exposed to Super RX-N films (Fujifilm) or revealed with iBright™ Imaging System (ThermoFisher). Protein expression was normalized to α-actinin or Gadph, or to total protein levels of the corresponding phosphorylated form. The list of antibodies is provided in Supplementary Materials.

### Polymerase chain reaction

Total RNAs were extracted with the RNeasy Mini Kit (Qiagen), reverse transcribed with the SuperScript III First-Strand Synthesis System (Invitrogen), and amplified with the Power SYBR Green Master Mix (Applied Biosystems) or the Hot FirePol EvaGreen qPCR Mix (Solis BioDyne). Expression of specific spliced or pan transcripts was analysed by end-point PCR and electrophoresis, or by quantitative PCR with the Step-One software and normalization to *Tbp* expression. The list of primers is provided in Table S1.

### Histology and immunofluorescence

Muscles were frozen in liquid nitrogen-cooled isopentane. Eight-micrometre muscle sections were stained with H&E and observed with an upright microscope (Olympus). For immunostaining, sections were fixed with 4% paraformaldehyde (PFA) or kept unfixed, then blocked in PBS, 3% BSA, incubated sequentially with primary and secondary fluorescent antibodies (Invitrogen, Jackson ImmunoResearch), mounted with Vectashield medium (Vector), and observed with Leica or Zeiss fluorescent microscopes. Quantification of fibre type and size was done as previously reported.^25^

### Staining of muscle bundles

To analyse NMJ organization, muscles were bathed *ex vivo* with α-bungarotoxin-Alexa555 (2 µg/ml - Invitrogen) for 30 min, before being washed and fixed with 4% PFA. Muscle bundles were cut, permeabilized in PBS, 1% Triton-X100, and blocked in PBS, 1% BSA, 0.1% Triton-X100. Bundles were then successively incubated with primary antibodies against Neurofilament and Synaptophysin (to stain pre-synaptic compartment), and the corresponding secondary antibodies (Invitrogen, Jackson ImmunoResearch). Images were recorded using a Leica confocal microscope.

### AChR turnover and recycling

AChR turnover was assessed by injecting α-bungarotoxin-Alexa647 and -Alexa555 (25 pmoles - Invitrogen) into TA/EDL muscles at days 1 and 10, respectively (5 and 14 days after nerve cut when combined with denervation), as previously done.^26^ AChR recycling was assessed by injecting α-bungarotoxin-Biotin (25 pmoles - Invitrogen) and saturating dose of streptavidin-Alexa647 (50 pmoles - Invitrogen) into TA/EDL muscles at day 1, followed by injection of streptavidin-Alexa555 (50 pmoles - Invitrogen) at days 2 or 4. Upon AChR internalization, streptavidin-Alexa647 is released from Btx-biotin, which remains bound to AChRs.^27^ Only recycled AChRs (*i.e.,* bound to free Btx-biotin) are labelled by streptavidin-Alexa555. We compared the population of recycled receptors (pixels positive for Alexa555) to the population of all receptors labelled with Btx-Biotin (pixels labelled with Alexa555 or Alexa647). For turnover and recycling quantification, images were recorded using Leica or Zeiss confocal microscopes. Pixel dominance (old *vs.* new or stable *vs.* recycled receptors) was calculated using Fiji and Matlab software. The assays could be applied to TA and EDL muscles, but not to *gastrocnemius* muscle because of its size and heterogeneity.

### Statistics

Quantitative data are displayed as mean ± SEM of independent samples, with n (number of individual experiments) ≥ 3. Statistical analysis of values was performed using unpaired Student’s t test or two-way ANOVA test with Tukey’s multiple comparisons test correction, with a 0.05 level of confidence accepted for statistical significance.

## Acknowledgements

We thank Prof. C.A. Thornton and Prof. M.S. Swanson for the generous gift of *HSA^LR^* and *Mbnl1^ΔE3/ΔE3^* mice, as well as Dr.D. Ham and M. Reischl for their help with the macros for fibre typing and AChR turnover. MHC antibodies, developed by H.M. Blau and S. Schiaffino, were obtained from the Developmental Studies Hybridoma Bank (University of Iowa, Iowa City, Iowa, USA). This work was supported by the Swiss Foundation for Research on Muscle Diseases and the Swiss National Science Foundation.

## Declaration of interests

The authors declare that they have no conflict of interest.

## Notes

### Competing Interest Statement

The authors have declared no competing interest.

### Summary of Updates

Errors in the text have been corrected Quality of the images in Figure 1 has been improved

## REFERENCES

1 Lin X, Miller JW, Mankodi A, Kanadia RN, Yuan Y, Moxley RT, et al. Failure of MBNL1-dependent post-natal splicing transitions in myotonic dystrophy. Hum Mol Genet 2006; 15(13):2087–2097.

2 Miller JW, Urbinati CR, Teng-Umnuay P, Stenberg MG, Byrne BJ, Thornton CA, et al. Recruitment of human muscleblind proteins to (CUG)(n) expansions associated with myotonic dystrophy. Embo J 2000; 19(17):4439–4448.

3 Nakamori M, Sobczak K, Puwanant A, Welle S, Eichinger K, Pandya S, et al. Splicing biomarkers of disease severity in myotonic dystrophy. Ann Neurol 2013; 74(6):862–872.

4 Suenaga K, Lee KY, Nakamori M, Tatsumi Y, Takahashi MP, Fujimura H, et al. Muscleblind-like 1 knockout mice reveal novel splicing defects in the myotonic dystrophy brain. PLoS One 2012; 7(3):e33218.

5 Sobczak K, Wheeler TM, Wang W, Thornton CA. RNA interference targeting CUG repeats in a mouse model of myotonic dystrophy. Mol Ther 2013; 21(2):380–387.

6 Martinez-Pena y Valenzuela I, Mouslim C, Akaaboune M. Calcium/calmodulin kinase II-dependent acetylcholine receptor cycling at the mammalian neuromuscular junction in vivo. J Neurosci 2010; 30(37):12455–12465.

7 Tang H, Macpherson P, Argetsinger LS, Cieslak D, Suhr ST, Carter-Su C, et al. CaM kinase II-dependent phosphorylation of myogenin contributes to activity-dependent suppression of nAChR gene expression in developing rat myotubes. Cell Signal 2004; 16(5):551–563.

8 Tang H, Macpherson P, Marvin M, Meadows E, Klein WH, Yang XJ, et al. A histone deacetylase 4/myogenin positive feedback loop coordinates denervation-dependent gene induction and suppression. Mol Biol Cell 2009; 20(4):1120–1131.

9 Macpherson P, Kostrominova T, Tang H, Goldman D. Protein kinase C and calcium/calmodulin-activated protein kinase II (CaMK II) suppress nicotinic acetylcholine receptor gene expression in mammalian muscle. A specific role for CaMK II in activity-dependent gene expression. J Biol Chem 2002; 277(18):15638–15646.

10 Macdermot V. The histology of the neuromuscular junction in dystrophia myotonica. Brain 1961; 84:75–84.

11 Allen DE, Johnson AG, Woolf AL. The intramuscular nerve endings in dystrophia myotonica--a biopsy study by vital staining and electron microscopy. 1969; 105(Pt 1):1–26.

12 Coers C, Telerman-Toppet N, Gerard JM. Terminal innervation ratio in neuromuscular disease. II. Disorders of lower motor neuron, peripheral nerve, and muscle. Arch Neurol 1973; 29(4):215–222.

13 Engel A, Jerusalem F, Tsujihata M, Gomez M. The neuromuscular junction in myopathies: a quantitative ultrastructural study. In: Recent Advances in Myology: Proceedings of the Third International Congress on Muscle Diseases, International Congress Series no 360. Edited by Bradley W, Gardner-Medwin D, Walton J. New York: Elsevier; 1975: 132–143.

14 Drachman DB, Fambrough DM. Are muscle fibers denervated in myotonic dystrophy? Arch Neurol 1976; 33(7):485–488.

15 Walton JN, Irving D, Tomlinson BE. Spinal cord limb motor neurons in dystrophia myotonica. J Neurol Sci 1977; 34(2):199–211.

16 Panaite PA, Kuntzer T, Gourdon G, Lobrinus JA, Barakat-Walter I. Functional and histopathological identification of the respiratory failure in a DMSXL transgenic mouse model of myotonic dystrophy. Dis Model Mech 2013; 6(3):622–631.

17 Spilker KA, Wang GJ, Tugizova MS, Shen K. Caenorhabditis elegans Muscleblind homolog mbl-1 functions in neurons to regulate synapse formation. Neural Dev 2012; 7:7.

18 Wheeler TM, Krym MC, Thornton CA. Ribonuclear foci at the neuromuscular junction in myotonic dystrophy type 1. Neuromuscul Disord 2007; 17(3):242–247.

19 Klinck R, Fourrier A, Thibault P, Toutant J, Durand M, Lapointe E, et al. RBFOX1 cooperates with MBNL1 to control splicing in muscle, including events altered in myotonic dystrophy type 1. PLoS One 2014; 9(9):e107324.

20 Mankodi A, Logigian E, Callahan L, McClain C, White R, Henderson D, et al. Myotonic dystrophy in transgenic mice expressing an expanded CUG repeat. Science 2000; 289(5485):1769–1773.

21 Kanadia RN, Johnstone KA, Mankodi A, Lungu C, Thornton CA, Esson D, et al. A muscleblind knockout model for myotonic dystrophy. Science 2003; 302(5652):1978–1980.

22 Castets P, Lin S, Rion N, Di Fulvio S, Romanino K, Guridi M, et al. Sustained activation of mTORC1 in skeletal muscle inhibits constitutive and starvation-induced autophagy and causes a severe, late-onset myopathy. Cell Metab 2013; 17(5):731–744.

23 Brockhoff M, Rion N, Chojnowska K, Wiktorowicz T, Eickhorst C, Erne B, et al. Targeting deregulated AMPK/mTORC1 pathways improves muscle function in myotonic dystrophy type I. J Clin Invest 2017; 127(2):549–563.

24 Dimauro I, Pearson T, Caporossi D, Jackson MJ. A simple protocol for the subcellular fractionation of skeletal muscle cells and tissue. BMC research notes 2012; 5:513.

25 Ham DJ, Borsch A, Lin S, Thurkauf M, Weihrauch M, Reinhard JR, et al. The neuromuscular junction is a focal point of mTORC1 signaling in sarcopenia. Nat Commun 2020; 11(1):4510.

26 Castets P, Rion N, Theodore M, Falcetta D, Lin S, Reischl M, et al. mTORC1 and PKB/Akt control the muscle response to denervation by regulating autophagy and HDAC4. Nat Commun 2019; 10(1):3187.

27 Bruneau E, Sutter D, Hume RI, Akaaboune M. Identification of nicotinic acetylcholine receptor recycling and its role in maintaining receptor density at the neuromuscular junction in vivo. J Neurosci 2005; 25(43):9949–9959.

28 Wang ET, Treacy D, Eichinger K, Struck A, Estabrook J, Olafson H, et al. Transcriptome alterations in myotonic dystrophy skeletal muscle and heart. Hum Mol Genet 2019; 28(8):1312–1321.

29 Strack S, Khan MM, Wild F, Rall A, Rudolf R. Turnover of acetylcholine receptors at the endplate revisited: novel insights into nerve-dependent behavior. J Muscle Res 2015; 36(6):517–524.

30 Wang AH, Yang XJ. Histone deacetylase 4 possesses intrinsic nuclear import and export signals. Mol Cell Biol 2001; 21(17):5992–6005.

31 Tintignac LA, Brenner HR, Ruegg MA. Mechanisms Regulating Neuromuscular Junction Development and Function and Causes of Muscle Wasting. Physiol Rev 2015; 95(3):809–852.

32 Cohen TJ, Waddell DS, Barrientos T, Lu Z, Feng G, Cox GA, et al. The histone deacetylase HDAC4 connects neural activity to muscle transcriptional reprogramming. J Biol Chem 2007; 282(46):33752–33759.

33 Moresi V, Williams AH, Meadows E, Flynn JM, Potthoff MJ, McAnally J, et al. Myogenin and class II HDACs control neurogenic muscle atrophy by inducing E3 ubiquitin ligases. Cell 2010; 143(1):35–45.

34 Akaaboune M, Culican SM, Turney SG, Lichtman JW. Rapid and reversible effects of activity on acetylcholine receptor density at the neuromuscular junction in vivo. Science 1999; 286(5439):503–507.

35 Strack S, Petersen Y, Wagner A, Roder IV, Albrizio M, Reischl M, et al. A Novel Labeling Approach Identifies Three Stability Levels of Acetylcholine Receptors in the Mouse Neuromuscular Junction In Vivo. PloS one 2011; 6(6):e20524.

36 Lee KY, Li M, Manchanda M, Batra R, Charizanis K, Mohan A, et al. Compound loss of muscleblind-like function in myotonic dystrophy. EMBO Mol Med 2013; 5(12):1887–1900.

37 Cohen TJ, Barrientos T, Hartman ZC, Garvey SM, Cox GA, Yao TP. The deacetylase HDAC4 controls myocyte enhancing factor-2-dependent structural gene expression in response to neural activity. FASEB J 2009; 23(1):99–106.

38 Mejat A, Ramond F, Bassel-Duby R, Khochbin S, Olson EN, Schaeffer L. Histone deacetylase 9 couples neuronal activity to muscle chromatin acetylation and gene expression. Nat Neurosci 2005; 8(3):313–321.

39 De Angelis L, Balasubramanian S, Berghella L. Akt-mediated phosphorylation controls the activity of the Y-box protein MSY3 in skeletal muscle. Skelet muscle 2015; 5:18.

40 Thomas JL, Moncollin V, Ravel-Chapuis A, Valente C, Corda D, Mejat A, et al. PAK1 and CtBP1 Regulate the Coupling of Neuronal Activity to Muscle Chromatin and Gene Expression. Mol Cell Biol 2015; 35(24):4110–4120.

