## Supplemental Data for "Destabilization of neuromuscular junctions and deregulation of activity-dependent signalling pathways in Myotonic Dystrophy type I"

Supplementary data includes: Supplementary Material, 8 Supplemental figures, 1 Supplementary Table.

### SUPPLEMENTARY MATERIAL

#### Antibodies

The following antibodies were used for immunoblotting (dilution 1/1000) or immunoflurescence: HDAC4 (#15164 and #7628; 1/5000 for IHC), Phospho-HDAC4Ser632 (#3424), Phospho-CaMKIIThr286 (#12716), CaMKII pan (#4436), Acetyl-Histone H3Lys9/Lys14 (#9677), Histone H3 (#9717), Akt (#9272), Phospho-Akt$^{Ser473}$ (#9271), S6 Ribosomal Protein (#2217), Phospho-S6 Ribosomal Protein$^{Ser235/6}$ (#2211), LC3B (#2775), GAPDH (#2118) from Cell Signaling Technology; α-actinin (A5044) and Neurofilament 200 (N4142; 1/2000 for IHC) from Sigma; Laminin (ab11575 and ab11576; 1/300 for IHC) from Abcam; Synaptophysin (A0010; 1/200 for IHC) from Dako.

**Fluorescence *in situ* hybridization**

FISH was conducted on muscle cryosections as previously described by Batra et al., [1] using a Cy3-CAG10 DNA probe. Nuclear foci were observed with a Leica confocal microscope.

**RNA-seq**

Obtained single-end RNA-seq reads were mapped to the mouse genome assembly (version mm10) with RNA-STAR (version 2.5.2a),[2] with default parameters except for allowing up 10 hits to genome (outFilterMultimapNmax 10), reporting only one location for hits with equal score (outSAMmultNmax 1), and for filtering reads without evidence in spliced junction table (outFilterType "BySJout"). All subsequent gene expression data analysis was done within the R software (R Foundation for Statistical Computing, Vienna, Austria). Raw reads and mapping quality was assessed by the qQCreport function from the R/Bioconductor software package QuasR (version 1.16.0).[3] Using RefSeq mRNA coordinates from UCSC (*genome.ucsc.edu*, downloaded in December 2015) and the qCount function from QuasR package, we quantified gene expression as the number of reads that started within any annotated exon of a gene. The differentially expressed genes were identified using the edgeR package (version 3.18.1).[4] The same RefSeq annotation was used to build the set of all existing junctions in transcripts, and qCount function was used to count the number of spanning reads in individual samples. The junctions were further filtered, only junction longer than 5 bp, and starting and ending in the same gene were kept, and those supported by at least 5 reads in total. The differentially used junctions were identified using the diffSpliceDGE function implemented in the edgeR package. Sequencing data have been deposited at the GEO repository and will be available upon publication.

### References


S1 Batra R, Charizanis K, Manchanda M, Mohan A, Li M, Finn DJ, et al. Loss of MBNL leads to disruption of developmentally regulated alternative polyadenylation in RNA-mediated disease. *Mol Cell* 2014; 56(2):311-322.

S2 Dobin A, Davis CA, Schlesinger F, Drenkow J, Zaleski C, Jha S, et al. STAR: ultrafast universal RNA-seq aligner. 2013; 29(1):15-21.

S3 Gaidatzis D, Lerch A, Hahne F, Stadler MB. QuasR: quantification and annotation of short reads in R. 2015; 31(7):1130-1132.

S4 Robinson MD, McCarthy DJ, Smyth GK. edgeR: a Bioconductor package for differential expression analysis of digital gene expression data. 2010; 26(1):139-140.

S5 Wang ET, Treacy D, Eichinger K, Struck A, Estabrook J, Olafson H, et al. Transcriptome alterations in myotonic dystrophy skeletal muscle and heart. *Hum Mol Genet* 2019; 28(8):1312-1321.

**Figure S1**

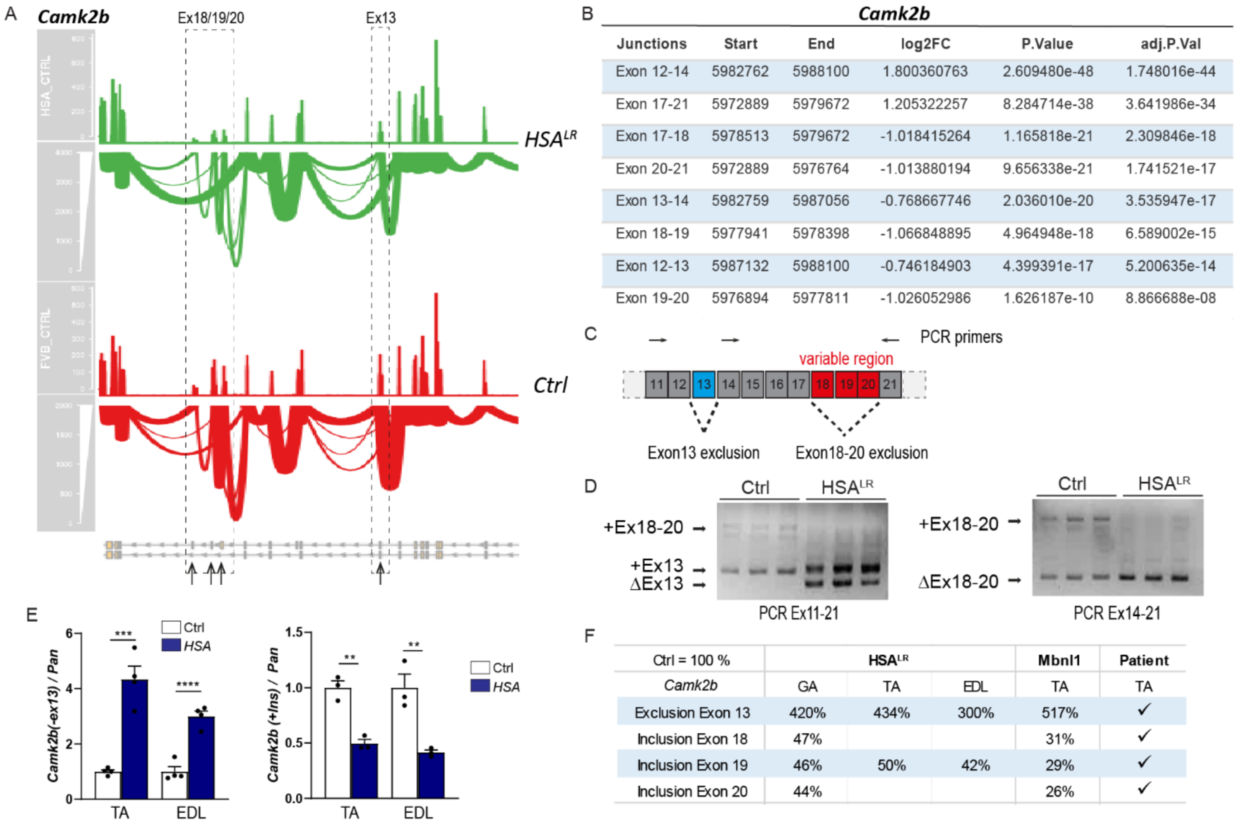


**Figure S1, related to Figure 2: Mis-splicing of *Camk2b* genes in *HSA^LR* muscle.** A, B) RNA-seq results for *Camk2b* showing mis-spliced regions in *HSA^LR* *gastrocnemius* muscle, compared to control. n=4. Statistical analysis is given in B, with the log2FC corresponding to the log-fold change in the expression of one exon, normalized to the expression of all the exons of the same gene, in *HSA^LR* muscle, as compared to controls. C, D) Splicing analysis of exon 13 and exons 18-20 of *Camk2b* by end-point PCR. Scheme of the gene and primers are given in C. E, F) Quantitative RT-PCR of the exclusion of exon 13 and of the inclusion of exons 18-20 in TA and EDL muscles from *HSA^LR* mice. Data are normalized on levels of total *Camk2b* transcripts. n = 4 (Ex13) and 3 (+Ins) per group. A summary table of *Camk2b* mis-splicing is given in F. All data are mean ± SEM; ** p<0.01; *** p<0.0001; **** p<0.0001; two-tailed unpaired Student’s t-test.

**Figure S2**

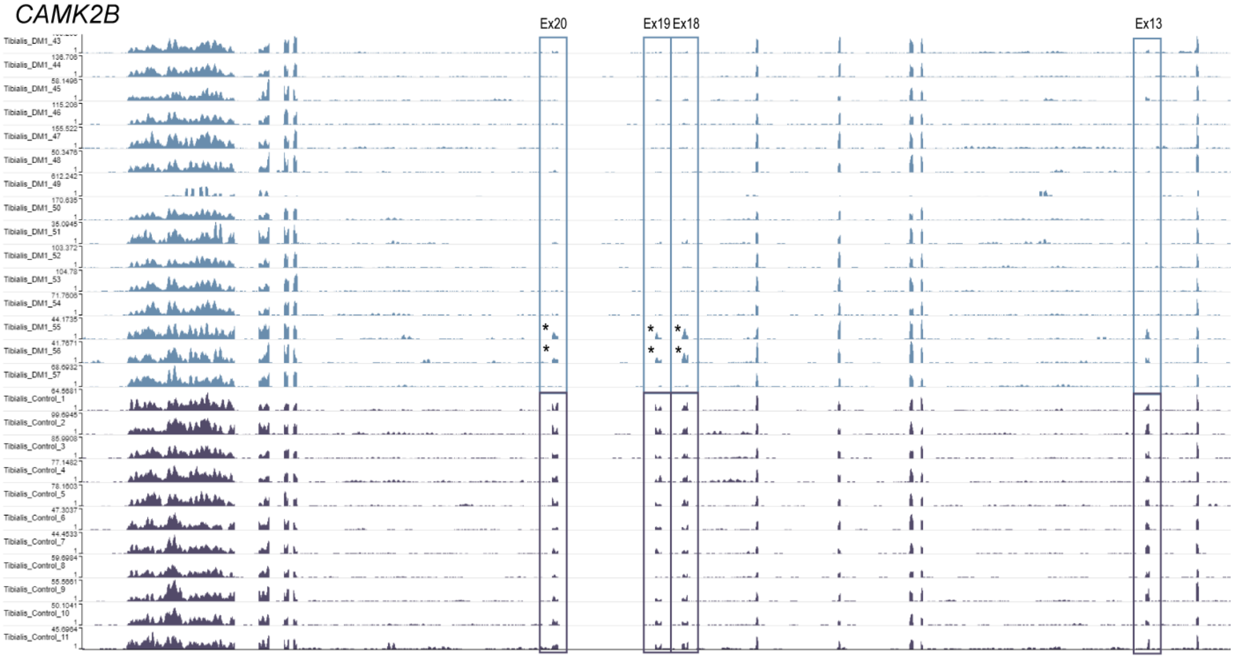


**Figure S2, related to Figure 2: Mis-splicing of *CAMK2B* gene in muscle from DM1 patients.** RNA-seq results for *CAMK2B* showing mis-splicing of exons 13, 18, 19 and 20 in TA muscle from DM1 patients, compared to control individuals. Data are from *dmseq.org.*[5] Asterisks show two DM1 muscles with normal splicing of *CAMK2B*.

**Figure S3**

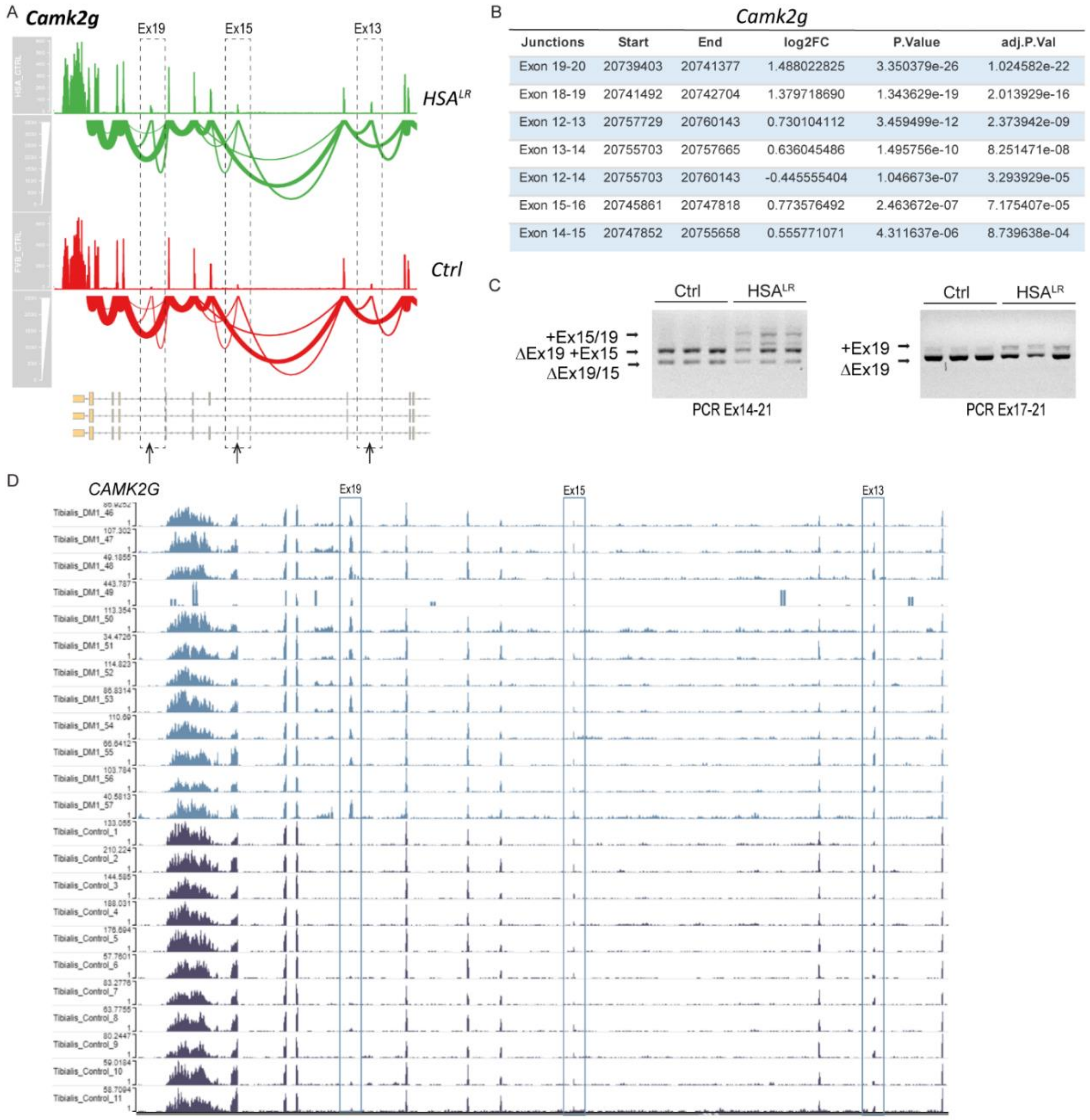


*Camk2g*

| Junctions | Start | End | log2FC | P.Value | adj.P.Val |
|---|---|---|---|---|---|
| Exon 19-20 | 20739403 | 20741377 | 1.488022825 | 3.350379e-26 | 1.024582e-22 |
| Exon 18-19 | 20741492 | 20742704 | 1.379718690 | 1.343629e-19 | 2.013929e-16 |
| Exon 12-13 | 20757729 | 20760143 | 0.730104112 | 3.459499e-12 | 2.373942e-09 |
| Exon 13-14 | 20755703 | 20757665 | 0.636045486 | 1.495756e-10 | 8.251471e-08 |
| Exon 12-14 | 20755703 | 20760143 | -0.445555404 | 1.046673e-07 | 3.293929e-05 |
| Exon 15-16 | 20745861 | 20747818 | 0.773576492 | 2.463672e-07 | 7.175407e-05 |
| Exon 14-15 | 20747852 | 20755658 | 0.555771071 | 4.311637e-06 | 8.739638e-04 |

**Figure S3, related to Figure 2: Mis-splicing of *Camk2g* genes in DM1 muscle.** A, B) RNA-seq results for *Camk2g* showing the mis-spliced regions in $HSA^{LR}$ muscle, compared to control. Statistical analysis is given in B, with the log2FC corresponding to the log-fold change in the expression of one exon, normalized to the expression of all the exons of the same gene, in $HSA^{LR}$ muscle, as compared to controls. n = 4 per group. C) End-point PCR analysis of mis-splicing of exons 15 and 19 of *Camk2g* in *gastrocnemius* from $HSA^{LR}$ and control muscles. n=3 per group. D) RNA-seq results for *CAMK2G* showing mis-splicing of exon 19 in TA muscle from DM1 patients, compared to control individuals. Data are from *dmseq.org.*[5]

**Figure S4**

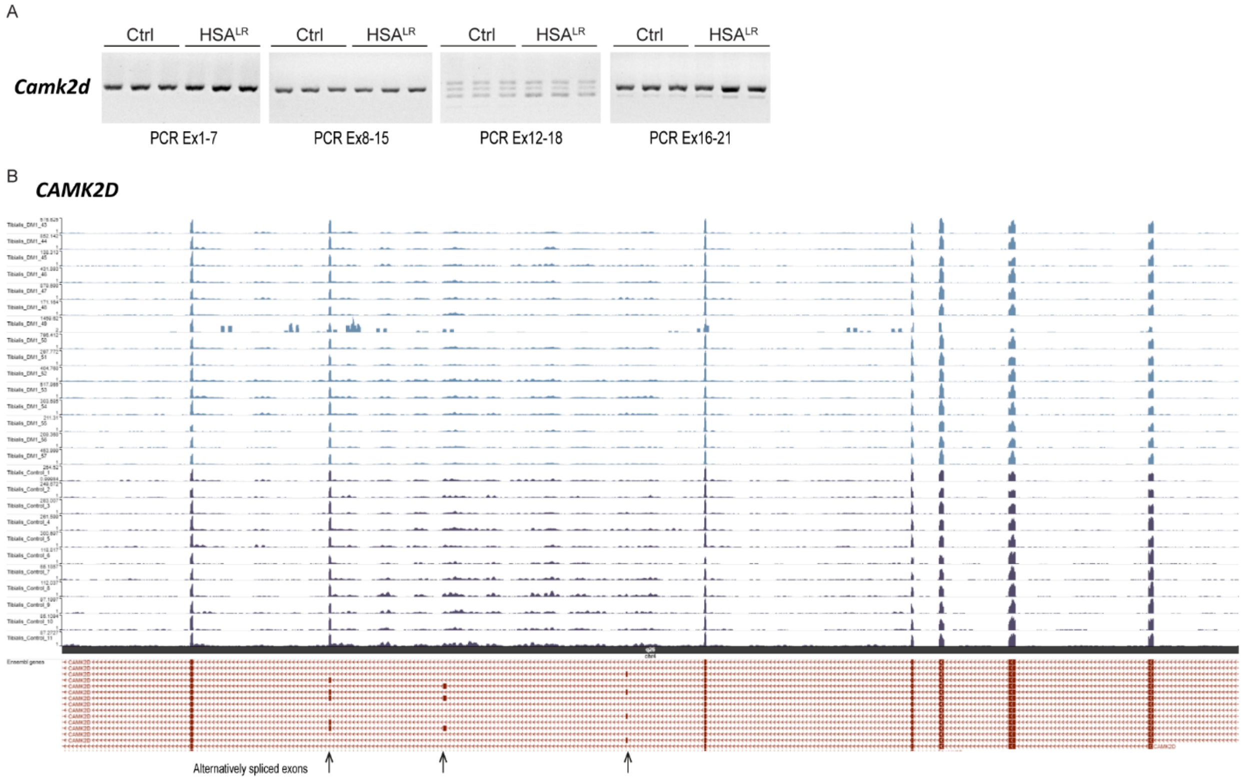


**Figure S4, related to Figure 2: Splicing of *CAMK2D* gene in muscle from DM1 patients.** A) End-point PCR analysis of exons 1 to 21 splicing of *Camk2d* in *gastrocnemius* from *HSA$^{LR}$* and control muscles. n=3 per group. B) RNA-seq results for *CAMK2D* showing normal splicing of 3 predicted alternative exons in TA muscle from DM1 patients, compared to control individuals. Data are from *dmseq.org.*[5]

**Figure S5**

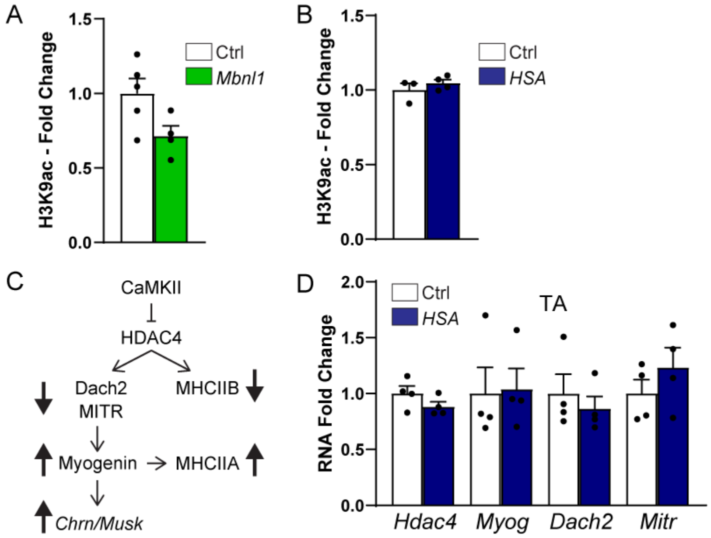


**Figure S5, related to Figure 5: Changes in HDAC4 downstream targets in *Mbnl1*$^{\Delta E3/\Delta E3}$ and *HSA*$^{LR}$ muscles.** A, B) Quantification of levels of acetylated histones (H3K9) in nuclear fraction of *gastrocnemius* muscle from *Mbnl1*$^{\Delta E3/\Delta E3}$ mice (A) and *HSA*$^{LR}$ mice (B). Levels are normalized to total histone H3 levels, and relative to control. n = 5/4 Ctrl/*Mbnl1*$^{\Delta E3/\Delta E3}$ (A) and 3/4 Ctrl/*HSA*$^{LR}$ (B). C) Regulation of activity-dependent pathways by CaMKII/HDAC4. D) Quantitative RT-PCR analysis of activity-dependent genes, *Hdac4, Myog, Dach2* and *Mitr*, in TA muscle from *HSA*$^{LR}$ mice. Transcript levels are normalized to *Tbp* and relative to control. n = 4 per group. All data are mean ± SEM.

**Figure S6**

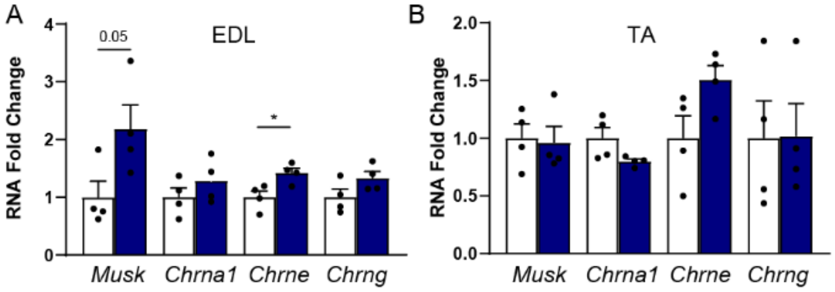


**Figure S6, related to Figure 6: Changes in synaptic gene expression in *Mbnl1$^{\Delta E3/\Delta E3}$* and *HSA$^{LR}$* muscles.** A, B) Quantitative PCR analysis of *Musk, Chrna1, Chrne* and *Chrng* in EDL (A) and TA (B) muscles from 3-month-old *HSA$^{LR}$* mice. Levels are normalized to *Tbp*. n=4 per group. All data are mean ± SEM; * $p<0.05$; two-tailed unpaired Student's t-test.

**Figure S7**

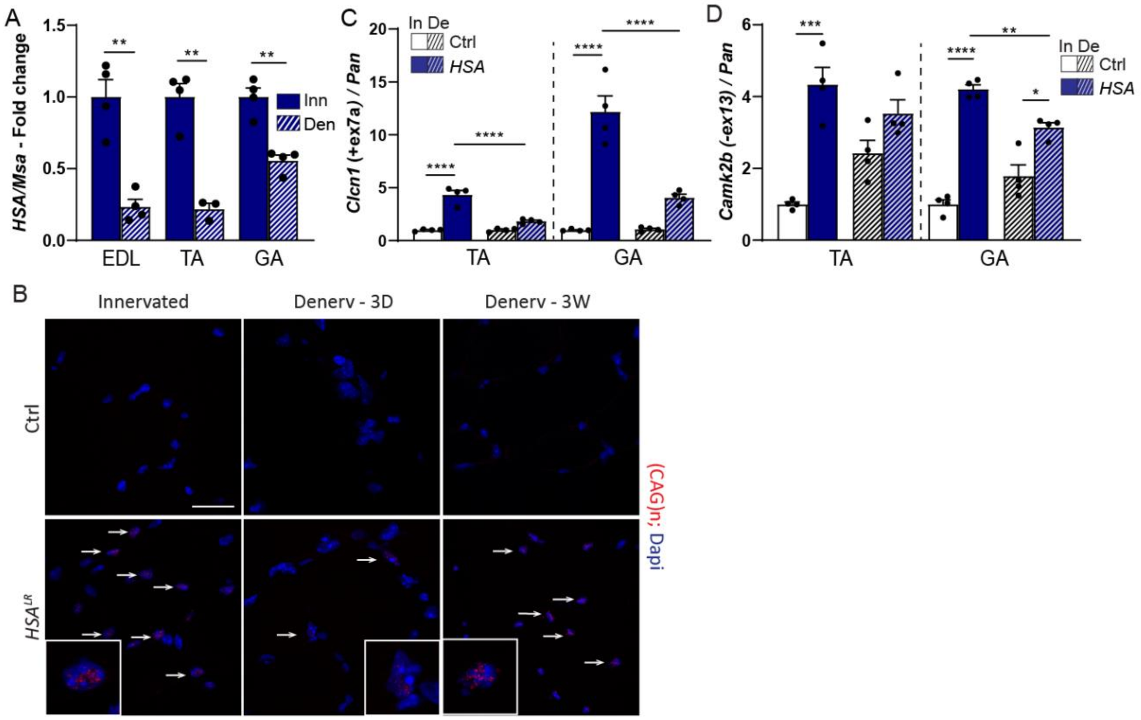


**Figure S7, related to Figure 7: Limits of the *HSA*$^{LR}$ mouse model upon nerve injury.** A) Quantitative PCR analysis of human *ACTA1* in EDL, TA and *gastrocnemius* in *HSA*$^{LR}$ innervated muscle and after 3 days of denervation. Results are normalized on mouse *Acta1* and relative to innervated *HSA*$^{LR}$ muscle. n=4 per group. B) Fluorescent *in situ* hybridization with Cy3-$CAG_{10}$ DNA probe, on sections of innervated and denervated (3 days and 3 weeks) muscles from control and *HSA*$^{LR}$ mice. Arrows point to positive nuclei. Scale bar, 25 µm. C, D) Quantitative PCR analysis of spliced transcripts of *Clcn1* (C) and *Camk2b* (D) in TA and *gastrocnemius* muscles from control and *HSA*$^{LR}$ mice after 3 days of denervation. Levels are normalized on total transcript levels and relative to innervated control muscle. n=4 per group. Data are mean ± SEM; *p<0.05; ** p<0.01; *** p<0.001; ****p<0.0001; two-way ANOVA with a Tukey's post-hoc analysis.

**Figure S8**

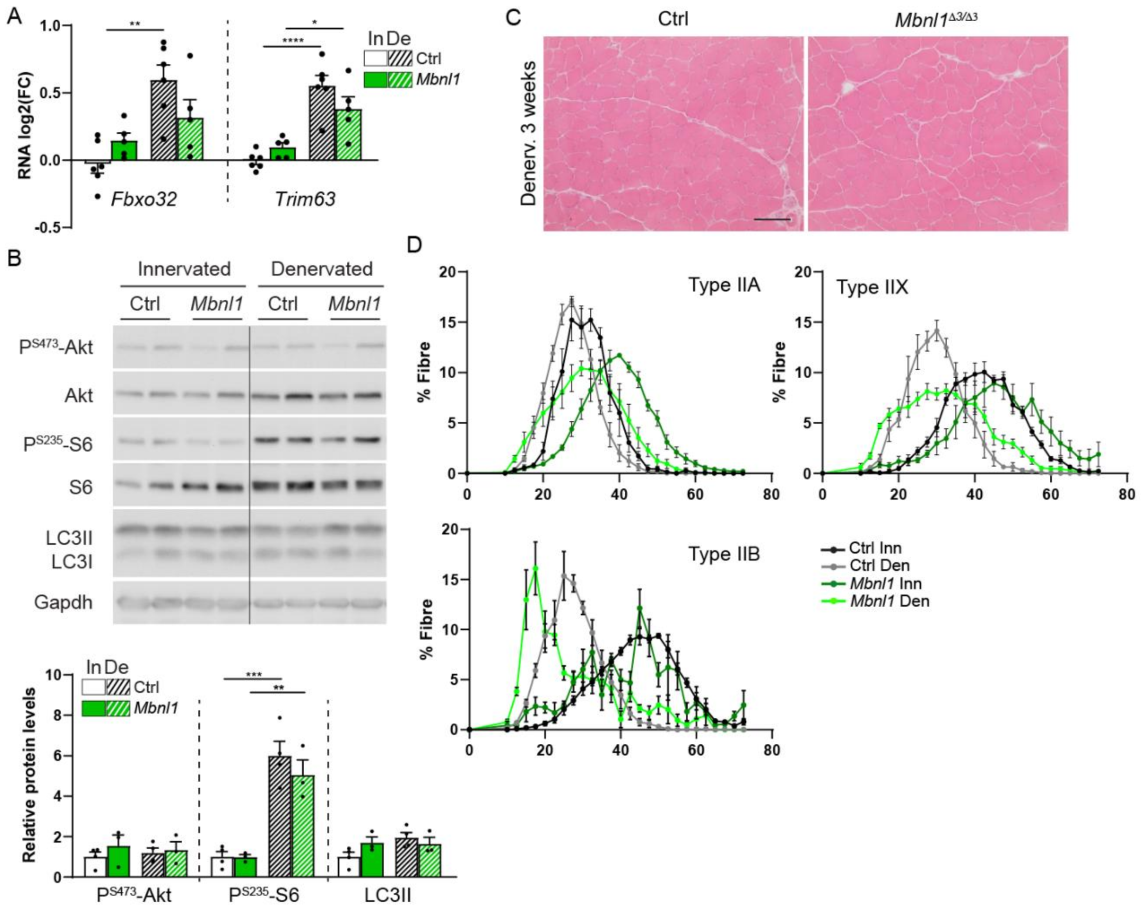


**Figure S8, related to Figure 7: Muscle response to denervation in *Mbnl1*$^{\Delta E3/\Delta E3}$ mice.** A) Quantitative PCR analysis of *Fbxo32* and *Trim63* in innervated and 3-day-denervated muscles from *Mbnl1*$^{\Delta E3/\Delta E3}$ mice. Transcript levels are normalized to *Tbp* and relative to control. n=6/5 Ctrl/*Mbnl1*$^{\Delta E3/\Delta E3}$. B) Western blot analysis of Akt/mTORC1 pathway in total protein lysate of TA muscle from *Mbnl1*$^{\Delta E3/\Delta E3}$ mice. Quantification of Akt/S6 phosphorylated levels and of LC3II is given in B. Levels are normalized to Gapdh, and relative to control. n = 4/3 Ctrl/*Mbnl1*$^{\Delta E3/\Delta E3}$. C) H&E coloration shows limited histopathologic alterations in denervated muscle from *Mbnl1*$^{\Delta E3/\Delta E3}$ and control mice, 3 weeks after nerve injury. Scale bar, 100 µm. D) Fibre size distribution for type IIA, IIX and IIB fibres in innervated and 3-week-denervated muscles from *Mbnl1*$^{\Delta E3/\Delta E3}$ and control mice. n = 4/4 Ctrl/*Mbnl1*$^{\Delta E3/\Delta E3}$. All data are mean ± SEM; * $p<0.05$; ** $p<0.01$; *** $p<0.001$; **** $p<0.0001$; two-way ANOVA with a Tukey's post-hoc analysis.

**Table S1: List of primers**

| Gene | Forward primer | Reverse primer |
|---|---|---|
| *Acta1 (mouse)* | 5'-CCGGAAAGAAATCTCAACCA-3' | 5'-CCAAGTCCTGCAAGTGAACA-3' |
| *ACTA1 (human)* | 5'-CGAGACCACCTACAACAGCA-3' | 5'-GGCATACAGGTCCTTCCTGA-3' |
| *Atp2a1+ex22* | 5'- GCCCTGGACTTTACCCAGTG-3' | 5'-ACGGTTCAAAGACATGGAGGA-3' |
| *Atp2a1 pan* | 5'- GCCCTGGACTTTACCCAGTG-3' | 5'-CCTCCAGATAGTTCCGAGCA-3' |
| *Camk2b-ex13* | 5'-TTTCTCAGCAGCCAAGAGTTT-3' | 5'-TTCCTTAATCCCGTCCACTG-3' |
| *Camk2b pan* | 5'-GCACGTCATTGGCGAGGA-3' | 5'-ACGGGTCTCTTCGGACTGG-3' |
| *Camk2b-ex18* | 5'-CCTGATGTCCTGAGCTTGGT-3' | 5'-GAACTGGAGATTGGCAGGAG-3' |
| *Camk2b-ex19* | 5'-TCAGTGAGAAGGGGCTGTG-3' | 5'-CTAGGAGACCCGGAGACAAG-3' |
| *Camk2b-ex20* | 5'-CCCCCAGGATCTCTGACA-3' | 5'-TGCTTCCGGGATGGGGTGGGC-3' |
| *Camk2b-ex13 (gel)* | 5'-GTTCCACCGTGGCCTCTAT-3' | 5'-TCGGAAGATTCCAGGGCAGC-3' |
| *Camk2b-ex18-20 (gel)* | 5'-CCAGACAAACAGCACCAAAA-3' | 5'-TGAGCTGCTCTGTGGTCTTG-3' |
| *Camk2g+exon 15/19 (gel)* | 5'-AGTTCCAGC GTGCACCTAAT-3' | 5'-ACGTGGACGTGAGGGTTTAG-3' |
| *Camk2g+exon 19 (gel)* | 5'-ACACCACTACAGAAGACGAAGA-3' | 5'-AACCTCAAACGAACAGGACC-3' |
| *Clcn1+ex7a* | 5'-GGGCGTGGGATGCTACTTTG-3' | 5'- AGGACACGGAACACAAAGGC-3' |
| *Clcn1 pan* | 5'-CTGACATCCTGACAGTGGGC-3' | 5'- AGGACACGGAACACAAAGGC-3' |
| *Chrna1* | 5'-TCCCTTCGATGAGCAGAACT-3' | 5'-GGGCAGCAGGAGTAGAACAC-3' |
| *Chrng* | 5'-GTGTCTTCGAGGTGGCTCTC-3' | 5'-TCTGGGATTGGAAGATGAGG-3' |
| *Dach2* | 5'-CCAGCTCAAATCCCAGTCAT-3' | 5'-CGCAGTTCCTTCTTTTCCTG-3' |
| *Hdac4* | 5'-CAGACAGCAAGCCCTCCTAC-3' | 5'-AGACCTGTGGTGAACCTTGG-3' |
| *Mitr* | 5'-CCTGCAGCACCTACTGTTGA-3' | 5'-GTACCTCTAATGCCCGGTGA-3' |
| *Myogenin* | 5'-ACTCCCTTACGTCCATCGTG-3' | 5'-CAGGACAGCCCCACTTAAAA-3' |
| *Myh2* | 5'-ACAAATCTATCCAAGTTCCG-3' | 5'-TTCGGTCATTCCACAGCATC-3' |
| *Myh4* | 5'-CAGATGAAAAGGTGGCCATT-3' | 5'-CTTCCCTTTGCTTTTGCTTG-3' |
| *Tbp* | 5'-CTCAGTTACAGGTGGCAGCA-3' | 5'-CAGCACAGAGCAAGCAACTC-3' |
